# Distinct metabolic flux modes through the tricarboxylic acid cycle in mesophyll and guard cells revealed by GC-MS-based ^13^C-positional isotopomer analysis

**DOI:** 10.1101/2021.02.16.431495

**Authors:** André G. Daubermann, Valéria F. Lima, Markus Schwarzländer, Alexander Erban, Joachim Kopka, Alisdair R. Fernie, Leticia dos Anjos, Danilo M. Daloso

**Affiliations:** Departamento de Biologia, Setor de Fisiologia Vegetal, Universidade Federal de Lavras, 37200-900, Lavras-MG, Brasil; LabPLant, Departamento de Bioquímica e Biologia Molecular, Universidade Federal do Ceará, 60451-970, Fortaleza-CE, Brasil; Westfälische Wilhelms-Universität Münster Institute of Plant Biology and Biotechnology, D-48143 Münster, Germany; Max-Planck-Institute of Molecular Plant Physiology, D-14476 Potsdam-Golm, Germany

**Author notes:** **Corresponding authors:** and, **Twitter account:** @danilodaloso; @LetsdosAnjos. These authors contributed equally (André G. Daubermann, Valéria F. Lima). **Brief heading:** MS-based ^13^C-positional labelling reveals specific guard cell metabolic flux modes through the tricarboxylic acid cycle and TRX regulation of fluxes from PEPc and PDH to Glu in illuminated mesophyll cells.

**Keywords:** ^13^C metabolic flux analysis, metabolic regulation, PEPc, PDH, mitochondrial thioredoxin, isotopomer analysis

## Abstract

- ^13^C-Metabolic flux analysis (^13^C-MFA) have greatly contributed to revealing the regulation of plant metabolism. However, mass spectrometry (MS) approaches have hitherto been limited in their power to deduce flux information due to lack of positional information.
- Here we established an MS-based ^13^C-positional isotopomer labelling approach and performed a multi-species/cell-types analysis based on previous ^13^C-MFA to compare flux modes through the tricarboxylic acid (TCA) cycle and associated pathways in mesophyll (MCs) and guard cells (GCs).
- Both cell types showed high ^13^C-enrichment in pyruvate. However, GCs and sink MCs, but not source MCs showed high ^13^C-incorporation into Glu/Gln following provision of ^13^C-sucrose. Only GCs showed higher ^13^C-enrichment in the carbon 1 atom of Gln, which is derived from PEPc-mediated CO_2_ fixation. Increased ^13^C-enrichment in the carbon 1 of Glu was also observed in both *trxo1* and *ntra ntrb* mutants, but not in wild type Arabidopsis plants, following provision of ^13^C-glucose.
- Our results suggest that the mitochondrial thioredoxin system restricts the fluxes from PEPc and glycolysis to Glu in illuminated MCs and reveal that fluxes throughout the TCA cycle of GCs resemble those of sink MCs but operate different non-cyclic flux modes to support Gln synthesis in the light.

## Introduction

In animals and in most microorganisms, respiration is responsible for most of the cellular energy transformed. In plants, this is not always the case due to their ability to photosynthesize in the light. Since both photosynthesis and respiration, as well as photorespiration, require tight, but conditional integration, plant energy metabolism needs to be particularly flexible (Møller *et al*., 2020). Furthermore, the regulation of plant respiration is light-dependent and inhibited at illumination (Tcherkez *et al*., 2012), in which the non-plastidial thioredoxin (TRX) system acts as a negative regulator of the metabolic fluxes toward the mitochondrial (photo)respiratory metabolism (Sevilla *et al*., 2020). This idea is supported by the finding that the mitochondrial TRX *o1* deactivates succinate dehydrogenase (SDH), fumarase (FUM) and mitochondrial lipoamide dehydrogenase (mtLPD) *in vitro* and that the *trxo1* mutant displays increased carbon fluxes toward the TCA cycle and the photorespiratory metabolism, as well as increased respiratory rates (Daloso *et al*., 2015b; Florez-Sarasa *et al*., 2019; Reinholdt *et al*., 2019; Nietzel *et al*., 2020). Phosphorylation of pyruvate dehydrogenase (PDH) by PDH kinase is an additional mechanism that negatively regulates the carbon fluxes toward the TCA cycle in the light (Tovar-Méndez *et al*., 2003). Despite considerable advances in the understanding of the regulatory mechanisms by posttranslational modifications, protein-protein interaction, and metabolic channelling (Zhang *et al*., 2017, 2018), how the metabolic fluxes through the TCA cycle are choreographed by the interplay of specific individual regulatory mechanisms is not understood.

Among the metabolic pathways that depend on carbon from respiratory intermediates, the synthesis of glutamate (Glu), glutamine (Gln) and the associated amino acids family draws carbon from the C6-branch (from citrate to 2-oxoglutarate (2-OG)) of the TCA cycle. This idea is based on the fact that genetic or pharmacological interference with citrate synthase (CS), isocitrate dehydrogenase (IDH) or 2-oxoglutarate dehydrogenase (OGDH) activities substantially alters the amount of Glu and Gln produced in leaves (Araujo *et al*., 2008; Sienkiewicz-Porzucek *et al*., 2010; Sulpice *et al*., 2010). However, given that the metabolic fluxes toward the TCA cycle are inhibited in the light (Tcherkez *et al*., 2005, 2009; Gauthier *et al*., 2010), the source of carbon for Glu and Gln synthesis remained unclear until recently. Critical insight has come from a diel course genome scale metabolic model which predicted that organic acids, especially citrate, stored in the previous night are an important source of carbon to feed the C6-branch of the TCA cycle in the light (Cheung *et al*., 2014). According to this model, and in agreement with several other studies, the plant TCA cycle does not always work as a cycle. Several non-cyclic flux modes have been demonstrated in leaves and seed embryos (Sweetlove *et al*., 2010), highlighting the flexibility of the TCA cycle in plants as well as the likely requirement for elaborate regulation.

Detailed information regarding the metabolic fluxes through the TCA cycle has been obtained by both mass spectrometry (MS) and nuclear magnetic resonance (NMR)-based ^13^C-metabolic flux analyses (MFA) (Zamboni *et al*., 2009; Heise *et al*., 2014). NMR enables the analysis of the ^13^C incorporation at specific position in a molecule, which has been a critical advantage over MS approaches. However, the sensitivity of NMR is much lower than that of MS, which limits the power of NMR to small metabolic networks (Ratcliffe & Shachar-Hill, 2006; Antoniewicz, 2013). Thus, the establishment of a MS-based platform to analyse positional ^13^C-isotopomer labelling opens the possibility to improve the resolution of MFA. The analysis of positional isotopomers (i.e. the differentiation of not only the mass difference but also the atomic level resolution of where the label resides at the atomic level), allows the discrimination of which pathways the synthesis of a particular metabolite or to the activation of a metabolic pathway. For instance, ^13^C-NMR has recently been used to investigate the source of carbon for Glu synthesis in *Helianthus annuus* L. leaves following provision of ^13^CO_2_ (Abadie *et al*., 2017). The results of this study indicate that there is incorporation of one carbon from the anaplerotic CO_2_ assimilation catalysed by phospho*enol*pyruvate carboxylase (PEPc) activity into the carbon 1 of the Glu backbone and that no carbon from glycolysis is incorporated into Glu following provision of short time ^13^CO_2_ labelling (Abadie *et al*., 2017). Intriguingly, this result is in contrast to those obtained from ^13^C-isotope labelling experiments in guard cells (GCs), which showed high ^13^C incorporation into Glu/Gln and TCA cycle metabolites following provision of ^13^C-sucrose and ^13^C-HCO_3_ in the light, respectively (Daloso *et al*., 2015a, 2016b; Robaina-Estévez *et al*., 2017; Medeiros *et al*., 2018). This suggests that the metabolic fluxes toward the TCA cycle and Glu/Gln synthesis are differentially regulated in mesophyll cells (MCs) and GCs under illumination and highlights the need for tissue or cell specific investigations.

Changes in guard cell metabolism are essential to regulate stomatal opening according to the prevailing environmental condition (Sussmilch *et al*., 2019; Franzisky *et al*., 2020). GCs typically have few chloroplasts and low chlorophyll content (Willmer & Fricker, 1996), giving rise to lower photosynthetic capacity than MCs (Gotow *et al*., 1988; Lawson *et al*., 2003). However, GCs have a comparatively high number of mitochondria and a high respiration rate (Willmer & Fricker, 1996; Araújo *et al*., 2011). However, an understanding of the differences in flux modes through the TCA cycle between GCs and MCs is still lacking, even though it would provide the metabolic underpinning of their fundamentally different functions in leaves (Daloso *et al*., 2017). Here we established a gas chromatography-mass spectrometry (GC-MS)-based ^13^C-positional isotopomer labelling approach to investigate the distribution of the metabolic fluxes derived from PEPc and glycolysis throughout the TCA cycle and associated pathways. Our approach started with the assembly of a fragmentation map of trimethylsilyl (TMS)-derivatized compounds with further validation by analysing positionally ^13^C-labelled standards using both GC-electron impact-*time of flight*-MS (GC-EI-TOF-MS) at nominal mass resolution and high-resolution GC-atmospheric pressure chemical ionization-TOF-MS (GC-APCI-TOF-MS). We next performed a multi-species/cell-types analysis, tracking the ^13^C-enrichment in specific isotopomers using previously published data from ^13^C-isotope labelling experiments carried out in MCs of Arabidopsis (MC-C3) and maize (MC-C4) plants and in GCs of Arabidopsis and tobacco (Szecowka *et al*., 2013; Daloso *et al*., 2015a; Arrivault *et al*., 2017; Dethloff *et al*., 2017; Medeiros *et al*., 2018). We further analysed previous published data from ^13^C-isotope labelling experiments carried out in plants impaired in the mitochondrial thioredoxin system (Daloso *et al*., 2015b). The results are discussed in the context of the feasibility of the ^13^C-positional isotopomer labelling approach and on the regulation of the TCA cycle and guard cell metabolism in the light.

## Material and methods Data set acquisition

The dataset analysed here comprised of 320 chromatograms of primary metabolites identified by GC-EI-TOF-MS from Arabidopsis rosettes, Arabidopsis and maize leaves and from guard cell-enriched epidermal fragments of Arabidopsis and tobacco (Szecowka *et al*., 2013; Daloso *et al*., 2015b,a; Arrivault *et al*., 2017; Dethloff *et al*., 2017; Medeiros *et al*., 2018) (see Table S1 for details).

### Establishment of the ^13^C-positional isotopomer labelling approach

In order to investigate the ^13^C-positional isotopomer labelling in metabolites of, or associated to, the TCA cycle, we first assessed the Golm Metabolome Database and studies involving TMS-derivatization (De Jongh *et al*., 1969; Petersson, 1972; Leimer *et al*., 1977; Antoniewicz *et al*., 2007; Abadie *et al*., 2017; Souza *et al*., 2018; Okahashi *et al*., 2019) to identify the chemical structure of the TMS-derivatized compounds of interest after electron impact (EI) fragmentation. After that, an *in silico* analysis was performed to identify the major well-known fragmentation patterns resulting from EI ionization using the software ChemBioDraw 12.0 (CambridgeSoft, Cambridge, MA, USA). Finally, a specific EI-based fragmentation map of TMS compounds was assembled, containing the estimated cleavage groups and resulting carbon skeletons for each metabolite of interest, such as pyruvate (Pyr), the TCA cycle intermediates malate (Mal), fumarate (Fum) and succinate (Succ) and the amino acids Ala, Asp, Glu and Gln (Fig. **S1**).

### Mass spectrometry analysis

In order to confirm the fragmentation simulation obtained by our *in silico* analysis, we compared our data with those obtained from Okahashi and collaborators, in which positionally labelled TMS-derivatized organic acids were analysed via GC-EI-MS (Okahashi *et al*., 2019), and with the fragmentation obtained by analysing positionally labelled standards via nominal mass resolution GC-EI-TOF-MS and high mass resolution GC-APCI-TOF-MS with previously established methods and settings (Kopka *et al*., 2017; Erban *et al*., 2020).

### Determination of the ^13^C-enrichment in metabolites associated to the TCA cycle

We determined the relative isotopomer abundance (%) of the isotopologues (M, M1, M2…Mn) and the relative metabolite content as relative to the internal standard, i.e. ribitol or sorbitol, and fresh weight. The total relative ^13^C-enrichment was calculated by the multiplication of the relative isotopomer abundance of the isotopologues by the number of ^13^C atoms incorporated in each isotopomer (i.e. M0*0+M1*1+M2*2…Mn*n) divided by the number of carbon of the fragment, as described previously (Lima *et al*., 2018). The total relative ^13^C-enrichment shows the amount of ^13^C incorporated from the labelled substrate into each metabolite fragment, while the relative isotopomer abundance analysis indicates the number of labelled carbons incorporated into each metabolite fragment. Given that the experiments had different labelling times, we also determined the ^13^C-enrichment rate, dividing the total relative ^13^C-enrichment obtained at the last time-point of each experiment by the ^13^C-enrichment at time zero and then normalizing the data by the experiment labelling time, in minutes. Although the ^13^C incorporation into metabolites is not linear and may differ between experiments and compounds, we did find this approach worth to use for the comparison between GCs and MCs.

### Estimation of positional ^13^C-enrichment in glutamate and malate

We further estimated the ^13^C-enrichment in the carbons 1 of Glu and 4 of Mal by using different fragments of these compounds according to a previous established mathematical framework, with modifications (Beylot *et al*., 1993). In the case of Glu, the ^13^C-enrichment in the carbon 1 was obtained by using the total ^13^C-enrichment of the fragments *m/z* (mass-to-charge ratio) 348 and *m/z* 246, that contain the carbons 1, 2, 3, 4 and 5 and 2, 3, 4 and 5, respectively. In the case of Mal, the estimation of the ^13^C-enrichment in the carbon 4 was obtained by using the total ^13^C-enrichment of the fragments *m/z* 245 (carbons 1, 2, 3-4), *m/z* 233 (carbons 2, 3-4) and *m/z* 265 (carbon 2). For this, the total ^13^C-enrichment of each fragment was normalized by the time 0 of each experiment, which corresponds to the natural ^13^C abundance found in each sample. The relative total ^13^C-enrichment of each fragment was then multiplied by the number of carbons present in each fragment. The positional ^13^C-enrichment was then obtained by subtracting the relative total ^13^C-enrichment of different fragments. For instance, the positional ^13^C-enrichment in the carbon 1 of Glu was obtained by subtracting the relative total ^13^C-enrichment of the fragment *m/z* 348 from those obtained in the fragment *m/z* 246. In the case of Mal, we first obtained the positional ^13^C-enrichment in the carbon 1 by subtracting the relative total ^13^C-enrichment of *m/z* 245 from *m/z* 233. Given that the fragment *m/z* 265 contains the carbon 2 of the Mal backbone (Okahashi *et al*., 2019), we then obtained the ^13^C-enrichment in the carbons 3 and 4 by subtracting the relative total ^13^C-enrichment of *m/z* 233 from *m/z* 265. Given that 66% of the *m/z* 117 corresponds to the fragment with the carbon 4, we then estimated the ^13^C-enrichment in the carbon 4 as being 66% of the ^13^C-enrichment found in the carbons 3 and 4.

### Statistical analysis

The relative total ^13^C-enrichment throughout time was compared to the time zero by ANOVA and Dunnet’s test (*P* < 0.05), which allow multiple comparison with a specific control (Dunnett, 1955). Exception is made for the data from MC-C4 experiment, which contains only one biological replicate and, thus, did not allow statistical comparison to other experiments. Punctual comparisons between GCs and MC-C3 and between source and sink MC-C3 were carried out by Student’s *t*-test (*P* < 0.05). The ^13^C-enrichment rate in TCA cycle and related metabolites of Arabidopsis leaves under different labelled substrates were compared using ANOVA and Tukey test (*P* < 0.05).

## Results

### Establishment of a GC-MS-based ^13^C-positional isotopomer labelling approach

Our *in silico* analysis showed that the fragmentation at the TMS-carboxyl group in the carbon 1 position of Glu and Gln result in the fragments *m/z* 117 and *m/z* 246 in Glu and *m/z* 117 and *m/z* 245 in Gln, whilst the fragment *m/z* 156 also contains the carbons 2, 3, 4 and 5 of the Glu backbone, but with an additional loss of a TMS+OH group (Fig. **S1**). Since the incorporation of ^13^C atoms in a fragment causes a shift in the *m/z* ratio observed in the mass spectra, we deduced that after ^13^C-labelling the Glu fragment *m/z* 117 should change to 118 and the fragments *m/z* 246 and *m/z* 156 should change from 246 to 250 and 156 to 160, respectively, according to the number of labelled carbons incorporated. Analysis of Glu standard labelled in the carbon 1 (1-^13^C-Glu) or 5 (5-^13^C-Glu) indicates that *m/z* 156 does not have the carbon 1 but contains the carbon 5 of the Glu backbone (Fig. **S2a**). This further suggests that the fragment *m/z* 117 of both Glu and Gln carries the carbon 1 position, which is derived from PEPc fixation (Fig. **1a,b**) (Abadie *et al*., 2017). However, mass spectral analysis of 1-^13^C-Glu standard indicate that this assumption should be taken with caution. In fact, the intensity of *m/z* 118 increased substantially in 1-^13^C-Glu and is 2.7-fold higher than naturally labelled Glu (Fig. **S2b**). However, by contrast to the observed in *m/z* 156 of 5-^13^C-Glu analysis, the intensity of *m/z* 118 did not overcome the values of the non-labelled ion (*m/z* 117), suggesting a possible influence of adjacent nominal mass fragments and respective isotopologues. Indeed, detailed mass spectral analysis highlights that the intensity of the ions *m/z* 114-118 are similar (Fig. **S2c**). Therefore, the *m/z* 117 ion can be an isotopomer of *m/z* 115 and *m/z* 116 peaks, which possibly explains the higher intensity of the *m/z* 117, compared to the intensity *m/z* 118 in 1-^13^C-Glu standard analysis (Fig. **S2b**). We then decided to calculate the ^13^C-enrichment in the carbon 1 of Glu by using information from the fragments *m/z* 348 and *m/z* 246. Interestingly, the relative total ^13^C-enrichment in the carbon 1 of Glu is linearly correlated with the relative total ^13^C-enrichment observed in the fragment *m/z* 117 (Fig. **S2d**). This reinforces the idea that this fragment contains the carbon 1 of the Glu backbone. Thereafter, we used the ^13^C-enrichment in *m/z* 246, *m/z* 245 and *m/z* 117 fragments to distinguish the relative contribution of glycolysis and TCA cycle pathways (*m/z* 246 and *m/z* 245) from that of the CO_2_ fixation via PEPc (*m/z* 117) to the synthesis of Glu and Gln.

**Fig. 1.**
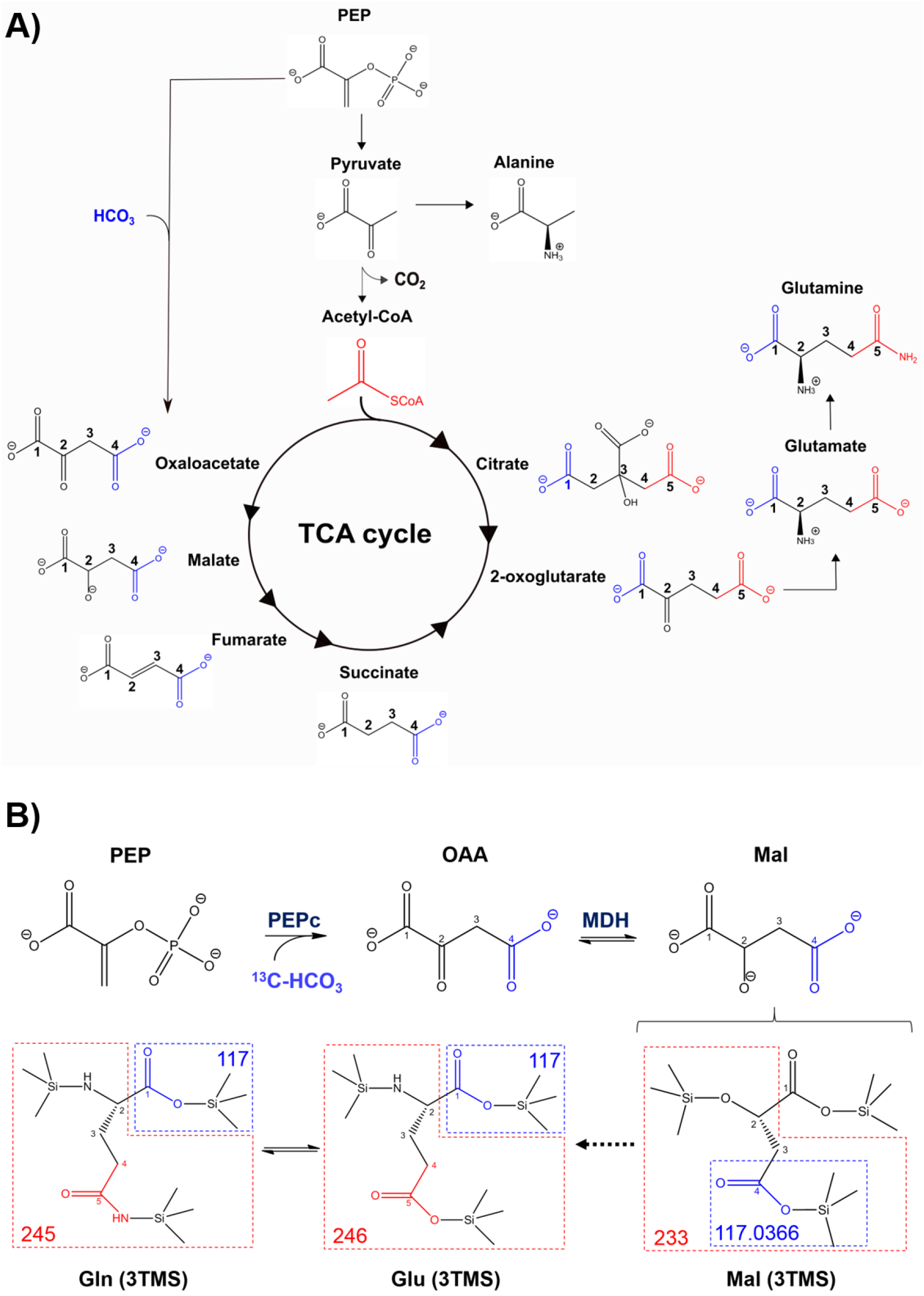
A) Simplified diagram of the TCA cycle and associated pathways, distinguishing the carbon fluxes from PEPc activity and glycolysis toward the TCA cycle and the synthesis of Glu and Gln. Carbons in blue and red types highlight the carbon derived from the PEPc-mediated anaplerotic CO_2_ fixation and those from Acetyl-CoA, respectively, according to a recent ^13^C-NMR metabolic flux study (Abadie *et al*., 2017). **B)** Upper part – Diagram showing the carbon tracking from PEPc-mediated ^13^C-HCO_3_ assimilation to the synthesis of OAA and Mal. Lower part – Schematic representation of TMS-derivatized Mal, Glu and Gln demonstrating the composition of representative fragments. Numbers correspond to the *m/z* ratio of each highlighted fragment. Abbreviations: Gln, glutamine; Glu, glutamate; Mal, malate; MDH, malate dehydrogenase; OAA, oxaloacetate; PEP, phospho*enol*pyruvate; PEPc, phospho*enol*pyruvate carboxylase.

The ^13^C-incorportion into the carbon 4 of Mal and/or Asp is an indirect evidence of PEPc activity (Abadie & Tcherkez, 2019). Analysis of positionally labelled organic acids by GC-EI-MS suggests that the Mal fragments *m/z* 117 and *m/z* 233 contain, respectively, the carbon 4 and the carbons 2, 3 and 4 of the Mal backbone (Okahashi *et al*., 2019). The *m/z* 117 is a common fragment obtained after EI-fragmentation of TCA cycle metabolites (see blue atoms in the Fig. **1a**), suggesting that this fragment could be used to investigate the distribution of the PEPc-derived carbon fixation throughout the TCA cycle (Fig. **1b**). However, our GC-APCI-MS analysis indicates the existence of two fragments with different monoisotopic masses, namely *m/z* 117,0366 and *m/z* 117,0730 (Fig. **S3,4**). These fragments correspond to 66 % and 34 % of the total *m/z* 117 abundance, respectively (Fig. **S4**). When comparing with 1-^13^C-Mal and 2-^13^C-Mal standards, it becomes clear that the fragment 117,0366 does not contain the carbons 1 and 2, whilst the fragment *m/z* 117,0730 contains the carbon 2 but not the 1 of the Mal backbone (Fig. **S4**). According to our fragmentation simulation and the data from Okahashi and collaborators, the fragment *m/z* 117,0366 contains carbon 4 of the Mal backbone (TMS-COO) whilst the fragment *m/z* 117,0730 possibly contains the carbons 2 and 3 (TMS-OCH2CH2) (Fig. **S1**). We next estimated the relative total ^13^C-enrichment in the carbon 4 of Mal, which was linearly correlated with the relative total ^13^C-enrichment in the fragment *m/z* 117 of Mal in Arabidopsis mesophyll cells under ^13^CO_2_, maize mesophyll cells under ^13^CO_2_ and guard cells under ^13^C-HCO_3_ (Fig. **S5**). Therefore, we propose that the PEPc-mediated carbon assimilation into Mal can be investigated by the analysis of *m/z* 117 obtained from EI fragmentation, but it is important to highlight that part of the ^13^C incorporation into this nominal mass fragment (less than 34%) does not correspond to the labelling incorporated by PEPc into the carbon 4.

Analysis of positional labelled standards by GC-APCI-MS and GC-EI-MS revealed which carbons of the Asp backbone are included in each major Asp fragment. This analysis showed that the fragment *m/z* 117 contains the carbons 3 and 4 and no other fragment contains only the carbon 4 of the Asp backbone (Fig. **S6**). Thus, in our approach, we used Mal, Glu and Gln, but not Asp, as an indirect measurement of the contribution of PEPc to fluxes throughout the TCA cycle and associated pathways. The power of this approach was revealed by performing a multi-species/cell-types analysis based on previous metabolic flux studies to compare flux modes through the TCA cycle and associated pathways in mesophyll (MCs) and guard cells (GCs) (Szecowka *et al*., 2013; Daloso *et al*., 2015b,a; Arrivault *et al*., 2017; Dethloff *et al*., 2017; Medeiros *et al*., 2018).

### Guard cells display lower carbon flux from pyruvate to alanine than mesophyll cells

No substantial changes in the content of Pyr, Ala and Asp were observed in MCs over time, but Asp and Pyr increased and decreased, respectively, in GCs (Fig. **S7**). The total relative ^13^C-enrichment in Pyr (*m/z* 174), Asp (*m/z* 232) and Ala (*m/z* 116) was higher in MC-C4 than in both MC-C3 and GCs (Fig. **2**). All carbon atoms of these metabolite fragments were labelled in both MC-C3 and MC-C4, as evidenced by the increases in the relative abundance of the isotopologues Pyr *m/z* 175-177, Asp *m/z* 233-235 and Ala *m/z* 117-118 (Fig. **S8**). However, MC-C4 displayed only slight increases in Asp *m/z* 118 (Fig. **2**; Fig. **S8**), while MC-C3 displayed a significant increase in the total relative ^13^C-enrichment in Asp *m/z* 117 (Fig. **2**). Pyr (*m/z* 174) was also rapidly labelled in GCs following provision of ^13^C-Suc. All carbon atoms of this fragment were labelled, as demonstrated by the increases in the relative abundance of the isotopologues Pyr *m/z* 175-177 (Fig. **S8**). However, no significant total relative ^13^C-enrichment in both Asp *m/z* 232 and *m/z* 117 was observed in GCs following provision of ^13^C-Suc. By contrast, an increase in total relative ^13^C-enrichment in either fragment of Asp was observed in GCs following provision of ^13^C-HCO_3_ (Fig. **2**), which was associated with slight increases in Asp *m/z* 118, *m/z* 234 and *m/z* 235 abundance (Fig. **S8**). GCs following provision of ^13^C-Suc showed a significant increase of total relative ^13^C-enrichment in Ala over time (Fig. **2**), although the increase in the abundance of the isotopologue Ala *m/z* 118 was less pronounced when compared to that observed in MCs (Fig. **S8**). Comparing the rates of relative total ^13^C-enrichment per minute, GCs displayed higher ^13^C-enrichment rate in Pyr and lower ^13^C-enrichment rate in Ala following provision of ^13^C-Suc, as compared to MC-C3 following provision of ^13^CO_2_ (Fig. **3**).

**Fig. 2.**
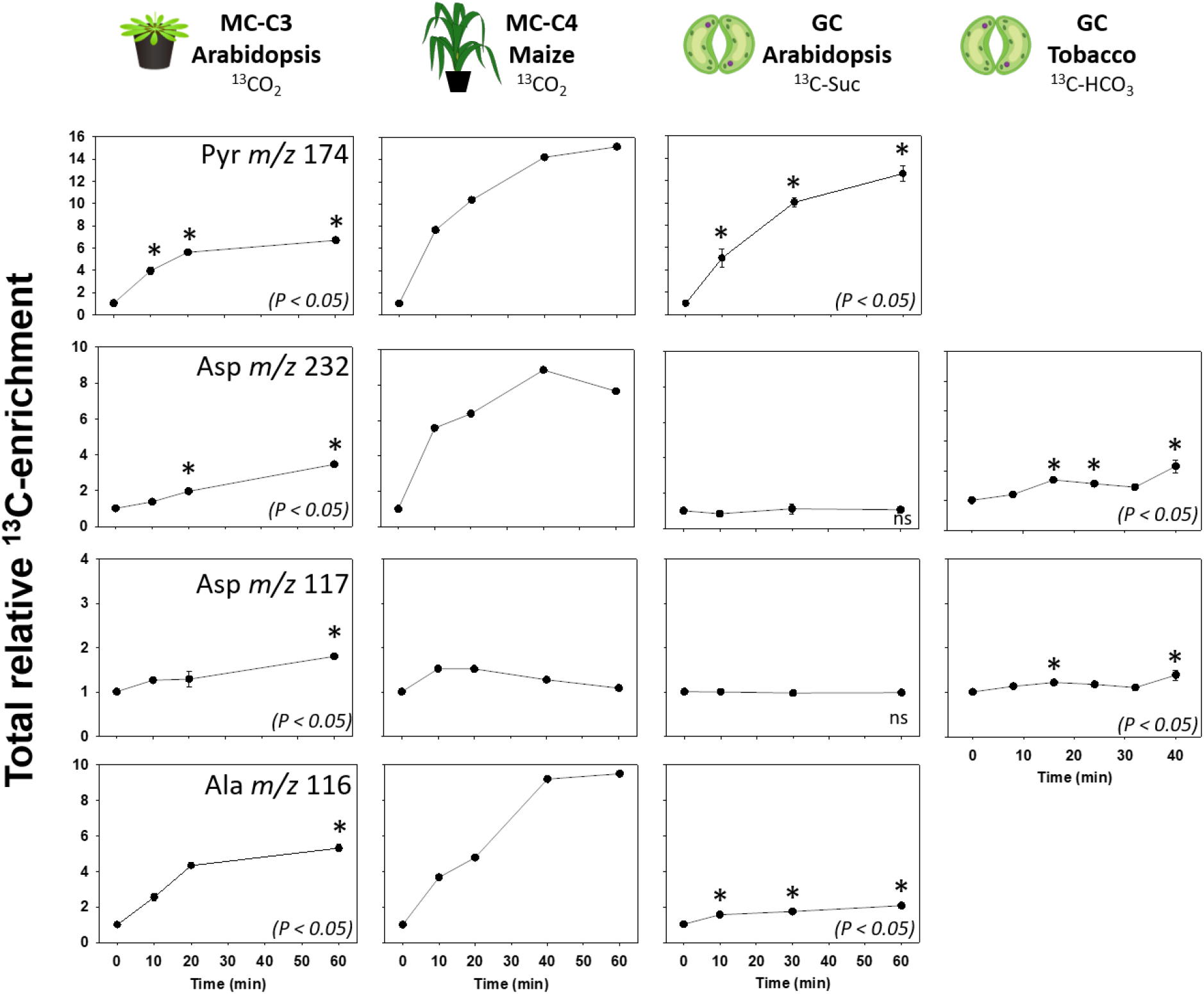
Total relative ^13^C-enrichment into Pyr (*m/z* 174), Asp (*m/z* 232 and 117) and Ala (*m/z* 116) from mesophyll cells (MCs) of Arabidopsis rosettes (MC-C3) and of maize plants (MC-C4) fed with ^13^CO_2_ for 60 minutes under light condition and from guard cells (GCs) of Arabidopsis rosettes fed with ^13^C-Suc for 60 minutes and of tobacco plants fed with ^13^C-HCO_3_ for 40 minutes, during dark-to-light transition. Total relative ^13^C-enrichment was calculated normalizing the total ^13^C-enrichment by the time zero. Values are presented as mean ± SE for MC-C3 (n=3) and GCs (n=4), except for MC-C4, in which biological replicates were absent. Asterisks indicate significant difference from time zero by Dunnet’s test (*P* < 0.05). Abbreviations: Ala, alanine; Asp, aspartate; Pyr, pyruvate.

### Guard cells show higher accumulation and ^13^C incorporation in fumarate than C3 mesophyll cells

The total relative ^13^C-enrichment in Mal, Fum and Succ was higher in MC-C4 than in MC-C3 and in GCs following provision of ^13^C-HCO_3_ (Fig. **4**) and absent in GCs following provision of ^13^C-Suc. The total relative ^13^C-enrichment in Fum (*m/z* 245) and Mal (*m/z* 117 and 233) increased in GCs and MCs over time. However, whilst GCs showed significant differences in the total relative ^13^C-enrichment of Fum after 16 min of labelling, MC-C3 exhibited significant difference in these metabolites only after 60 min. Additionally, Fum labelling was highly stable in MC-C3 over time, with slight increases in the abundance of the isotopologues Fum *m/z* 248 and *m/z* 249 at 60 min (Fig. **S9**). Increases in Fum *m/z* 249 indicate that all carbon atoms of this fragment were labelled in both MCs following provision of ^13^CO_2_ and in GCs following provision of ^13^C-HCO_3_ (Fig. **S9**). Interestingly, GCs displayed a higher decrease in the relative abundance of Fum *m/z* 245 than in Mal *m/z* 233 at the end of the experiment, while the opposite was observed in MC-C3 (Fig. **S9**). Furthermore, Fum content increased only in GCs (Fig. **S10**). When the time of labelling was considered, GCs following provision of ^13^C-HCO_3_ displayed higher ^13^C-enrichment rate in Fum than MC-C3 following provision of ^13^CO_2_ (Fig. **3**).

**Fig. 3.**
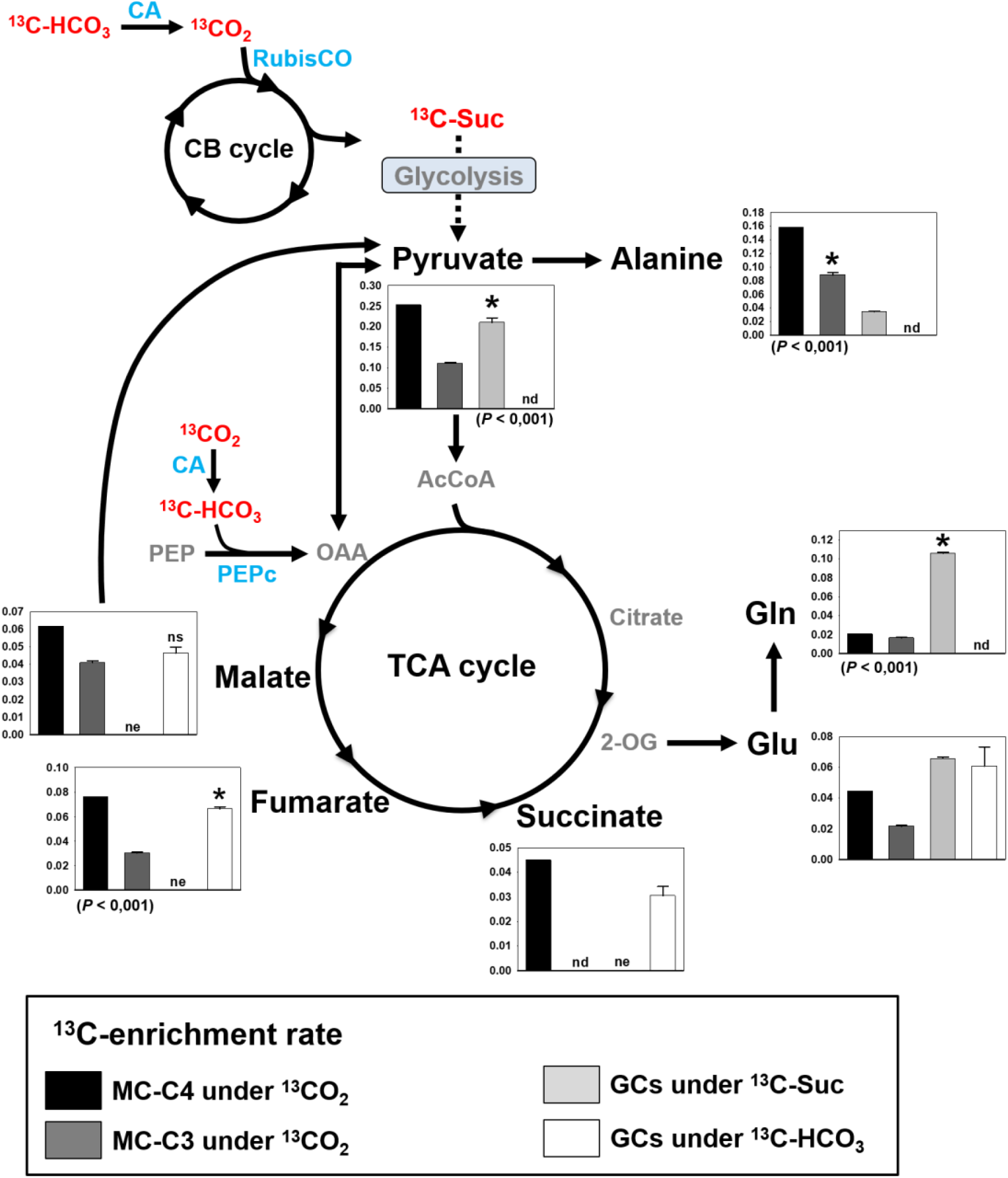
^13^C-enrichment rate of metabolites of, or associated to, the TCA cycle. The ^13^C-enrichment rate was determined by dividing the total relative ^13^C-enrichment by the ^13^C-feeding time, in minutes, of each cell type. Mesophyll cells (MCs) of Arabidopsis rosettes (MC-C3) and of maize plants (MC-C4) were fed with ^13^CO_2_ for 60 minutes under light condition. Guard cells (GCs) of Arabidopsis rosettes were fed with ^13^C-Suc for 60 minutes and of GCs of tobacco plants were fed with ^13^C-HCO_3_ for 40 minutes, during dark-to-light transition. Values are presented as mean ± SE for MC-C3 (n=3) and GCs (n=4), except for MC-C4, in which biological replicates were absent and, for this reason, statistical comparison with MC-C3 and GCs is also absent. Asterisks indicate significant difference between MC-C3 and GCs by Student’s *t*-test (*P* < 0.05). Metabolites that were not detected by GC-MS were identified as not analysed (nd). Metabolites with absent ^13^C-enrichment were identified as no ^13^C-enrichment observed (ne). Abbreviations: Metabolites: AcCoa, acetyl-coenzyme A; Glu, glutamate; Gln, glutamine; OAA, oxaloacetate; PEP, phospho*enol*pyruvate; 2-OG, 2-oxoglutarate. Enzymes, highlighted in blue: CA, carbonic anhydrase; PEPc, phospho*enol*pyruvate carboxylase; RubisCO, ribulose-1,5-biphosphate carboxylase/oxygenase.

MC-C4 had the highest ^13^C incorporation into the fragments *m/z* 117 and *m/z* 233 (Fig. **4**,**5**) and in the carbon 4 of Mal (Fig. **S11**). MC-C3 and GCs showed lower, but significant and nearly linear increases in Mal fragments over time. The ^13^C-enrichment rate in Mal *m/z* 117 and the ^13^C-incorporation into the carbon 4 was higher in MC-C3 than in GCs following provision of ^13^C-HCO_3_. However, no statistical difference in the rate of ^13^C-enrichment in *m/z* 233 was observed between these cell types (Fig. **5**). Interestingly, no total relative ^13^C-enrichment over time was observed in both Fum *m/z* 117 and Succ *m/z* 117 in GCs (Fig. **4**). Increased labelling in Succ *m/z* 247 was detected in GCs following provision of ^13^C-HCO_3_, reaching maxima of 17% at 32 minutes, when then it decreased towards 40 min (Fig. **4**). Similarly, Succ *m/z* 247 content increased linearly until 24 min and then decreased towards 40 min in GCs following provision of ^13^C-HCO_3_ (Fig. **S10**). By contrast to GCs, no substantial change in the content of Fum, Mal and Succ was observed in MCs over time (Fig. **S10**).

**Fig. 4.**
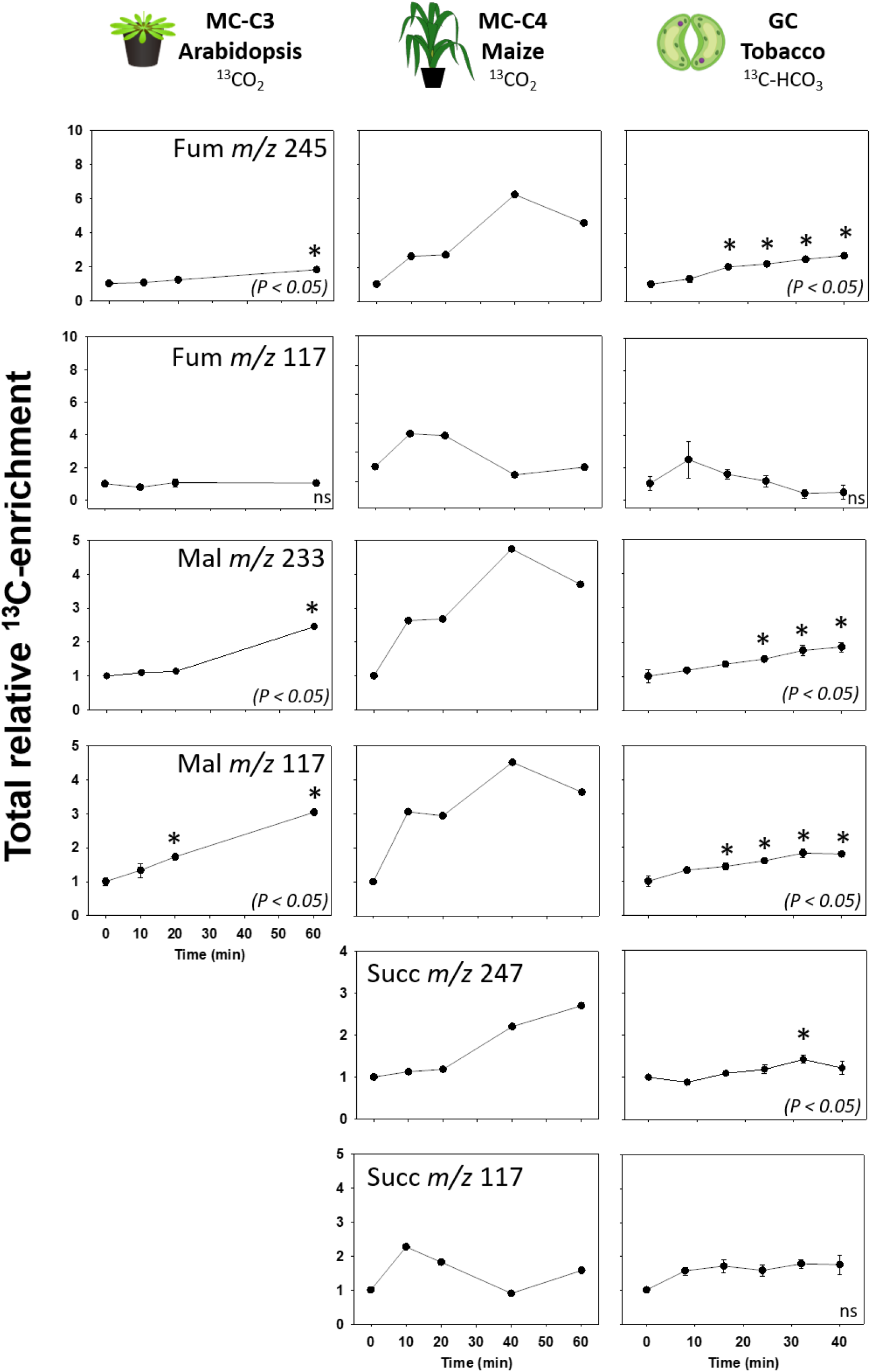
Total relative ^13^C-enrichment into the TCA cycle C4-branch metabolites from mesophyll cells (MCs) of Arabidopsis rosettes (MC-C3) and maize plants (MC-C4) and guard cells (GCs) of tobacco plants. The labelling time, the labelled substrates used, and the calculation of ^13^C-enrichment were carried out as described in the Figure 2. Asterisks indicate significant difference from time zero by Dunnet’s test (*P* < 0.05). Abbreviations: Fum, fumarate; Mal, malate; Succ, succinate.

**Fig. 5.**
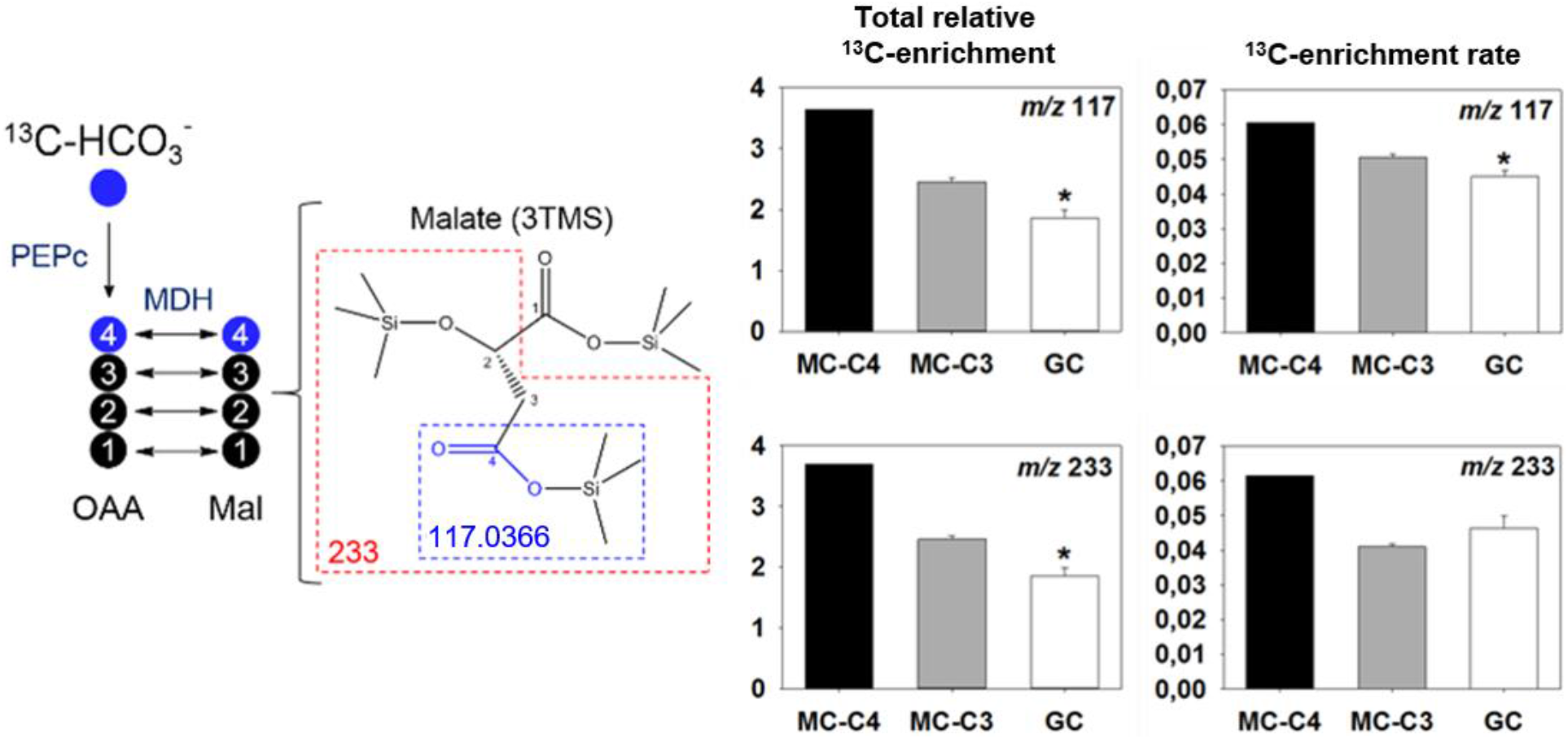
Schematic representation highlighting the reactions catalyzed by both PEPc and MDH, the fragments *m/z* 233 and *m/z* 117 of malate 3 TMS and the total relative ^13^C-enrichment and the ^13^C-enrichment rate obtained from both malate isotopomers (*m/z* 117 and *m/z* 233). The graphs refer to data from mesophyll cells (MCs) of Arabidopsis rosettes (MC-C3) and of maize plants (MC-C4) fed with ^13^CO_2_ for 60 minutes under light condition, and from guard cells (GCs) of tobacco plants fed with ^13^C-HCO_3_ for 40 minutes, during dark-to-light transition. The ^13^C-enrichment rate was determined by dividing the total relative ^13^C-enrichment by the ^13^C-feeding time, in minutes. Asterisks indicate significant difference between MC-C3 and GCs by Student’s *t*-test (*P* < 0.05). Abbreviations: Mal, malate; malate; MDH, malate dehydrogenase; OAA, oxaloacetate; PEPc, phospho*enol*pyruvate carboxylase.

### Guard cells show enhanced activation of the C6-branch of the TCA cycle toward glutamine synthesis as compared to mesophyll cells

GCs following provision of ^13^C-Suc had the highest total relative ^13^C-enrichment in Glu *m/z* 246 and Gln *m/z* 245 (Fig. **6**), which is related to a high increase in Glu *m/z* 248 and Gln 248 and 249 (Fig. **S12**). The total relative ^13^C-enrichment in Glu *m/z* 246 observed in GCs following provision of ^13^C-HCO_3_ resembled that observed in MC-C4 following provision of ^13^CO_2_, in which a slight increase in both Glu *m/z* 249 and Glu *m/z* 250 from 32 to 40 min was observed (Fig. **S12**). By contrast, MC-C3 showed a small increase over time of total relative ^13^C-enrichment in Glu *m/z* 246 and absent ^13^C-enrichment in Gln *m/z* 245 (Fig. **6)**. Additionally, GCs displayed significantly higher ^13^C-enrichment rate in Gln than both MC-C3 and MC-C4 (Fig. **3**). Regarding carbon 1 of Glu and Gln, which is derived from PEPc activity, GCs showed notable increases of 37 % and 26 % in the total relative ^13^C-enrichment of Glu *m/z* 117 and Gln *m/z* 117, respectively, while MC-C3 and MC-C4 showed minor changes over time (Fig. **6**; Fig. **S12**). Beyond being labelled mainly in GCs, the content of both Glu and Gln increased only in GCs following provision of ^13^C-Suc. Whilst Glu showed an accumulation peak at 10 min, followed by a decrease in its content, Gln showed a progressive and pronounced increase over time (Fig. **S13**). Interestingly, the accumulation and the ^13^C-incorporation in Glu coincide with the decrease in labelling and content observed in Succ in guard cells fed with ^13^C-HCO_3_ (Fig. **S14**).

**Fig. 6.**
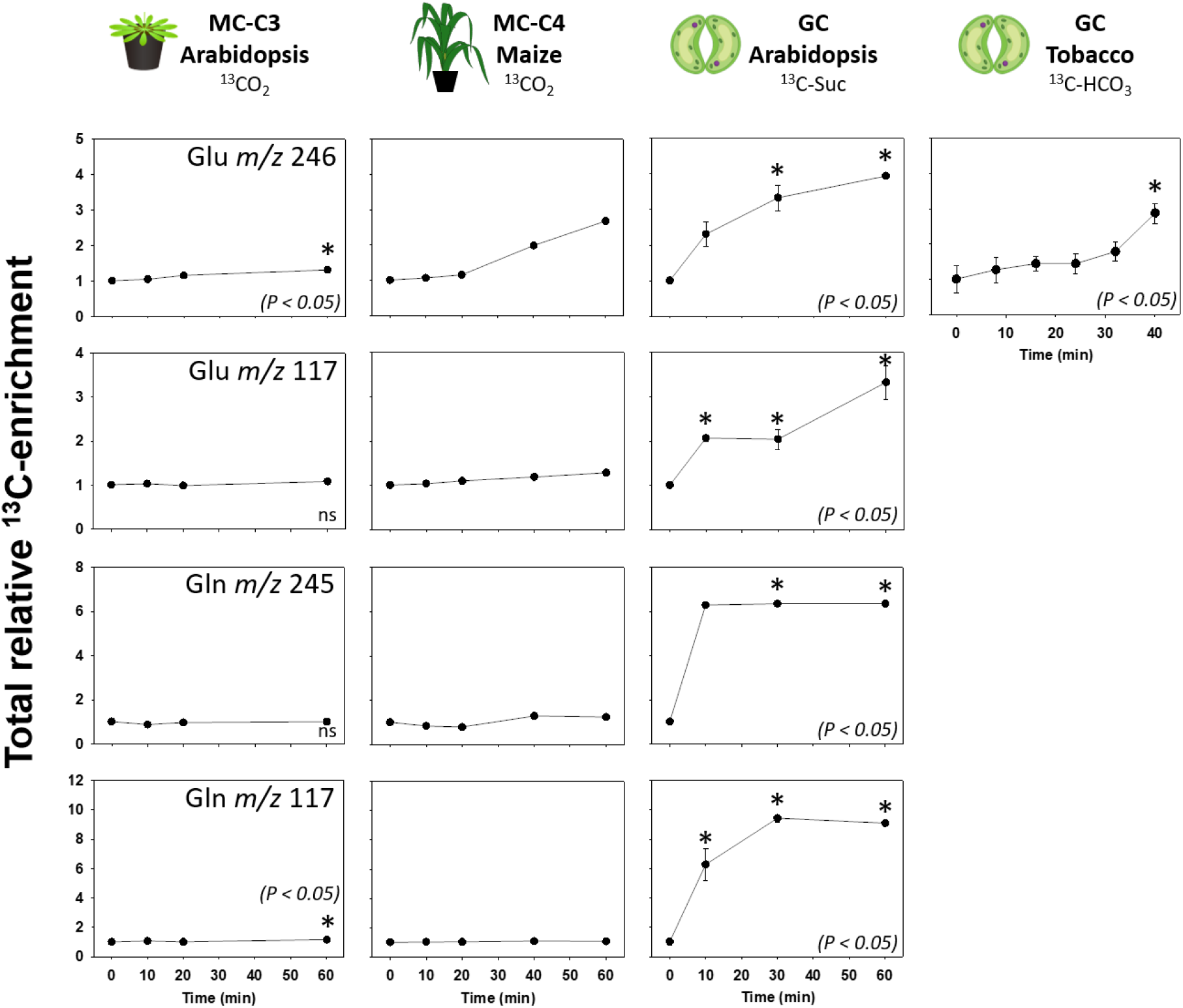
Total relative ^13^C-enrichment into Glu and Gln from mesophyll cells (MCs) of Arabidopsis rosettes (MC-C3) and maize plants (MC-C4) and guard cells (GCs) of tobacco plants. The labelling time, the labelled substrates used, and the calculation of ^13^C-enrichment were carried out as described in the Figure 2. Asterisks indicate significant difference from time zero by Dunnet’s test (*P* < 0.05). Abbreviations: Glu, glutamate; Gln, glutamine.

### Carbon flux to glutamate and glutamine is higher in sink than in source C3 mesophyll cells

We further analysed chromatograms of sink and source MCs of Arabidopsis plants fed with ^13^C-Suc for 240 min (Dethloff *et al*., 2017). The aim was to investigate differences between sink and source MC-C3 and to make a qualitative comparison of these results with those obtained from GCs following provision of ^13^C-Suc. Sink MC-C3 showed higher total relative ^13^C-enrichment in Mal, Fum and Succ than source MC-C3, although the levels of these metabolites were not different between these cell types (Fig. **7**). No statistical difference was found for total relative ^13^C-enrichment and relative metabolite content of Ala when comparing source and sink MC-C3. By contrast, the total relative ^13^C-enrichment and relative metabolite content of both Glu and Gln were higher in sink MC-C3. Interestingly, sink MC-C3 had higher maltose content, but no ^13^C-enrichment in this metabolite (Fig. **7**).

**Fig. 7.**
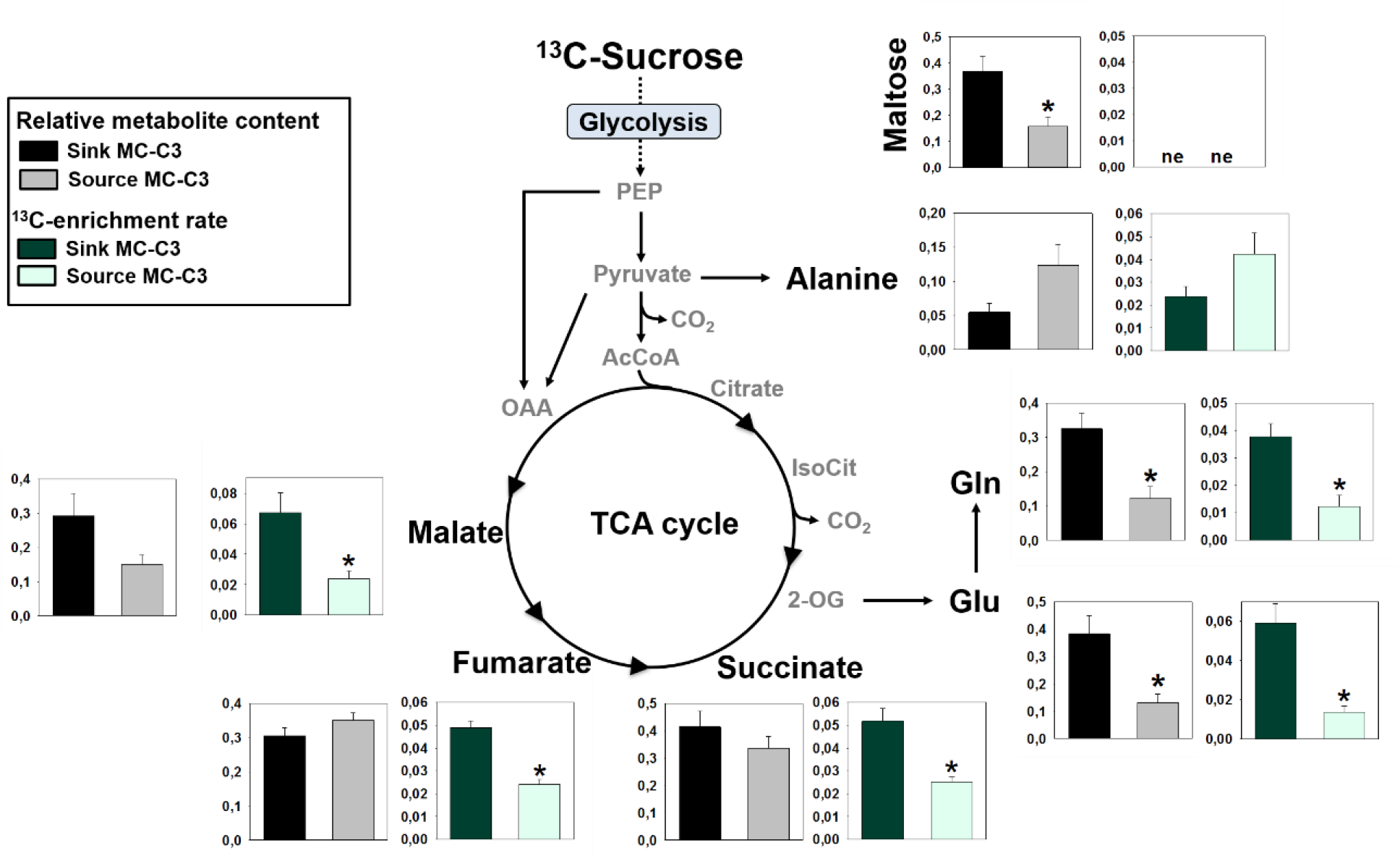
Diagram of TCA cycle and related metabolites showing relative metabolite content and ^13^C-enrichment rate of source and sink mesophyll cells (MCs) from Arabidopsis leaves (MC-C3) fed with ^13^C-Suc for 240 minutes, under light condition. The relative metabolite content was calculated in relation to ribitol and fresh weight (g FW^-1^) and the ^13^C-enrichment rate was determined by dividing the relative total ^13^C-enrichment by the ^13^C-feeding time, in minutes. Metabolite with absent ^13^C-enrichment is identified as no ^13^C-enrichment observed (ne). Values are presented as mean ± SE (n=6). Asterisks indicate significant difference between sink and source leaves by Student’s *t*-test (*P* < 0.05). Abbreviations: AcCoA, acetyl-coenzyme A; Glu, glutamate; Gln, glutamine; IsoCit, isocitrate; OAA, oxaloacetate; PEP, phospho*enol*pyruvate; Suc, sucrose; 2-OG, 2-oxoglutarate.

### PEPc-mediated incorporation into the carbon 1 of the glutamate backbone is negatively regulated by the mitochondrial NTR/TRX system

We assessed chromatograms of ^13^C-feeding experiments carried out in leaves of Arabidopsis wild type (WT) fed with ^13^C-glucose (Glc), ^13^C-Mal or ^13^C-Pyr (Daloso *et al*., 2015b). Following the provision of ^13^C-Mal ^13^C-enrichment in Pyr *m/z* 174 was observed, and the rate of the enrichment was even greater following provision of ^13^C-Glc (Fig. **8**). Fum showed higher ^13^C-enrichment rate following provision of ^13^C-Mal than following provision of ^13^C-Glc or ^13^C-Pyr. However, no difference was observed in the ^13^C-enrichment rate in both Mal and Fum when comparing ^13^C Glc and ^13^C-Pyr treatments. Furthermore, the rate of ^13^C-enrichment in Glu *m/z* 246 was highest following provision of ^13^C-Pyr. Lastly, ^13^C-enrichment in Glu *m/z* 117 was only observed following provision of ^13^C-Pyr (Fig. **8**).

**Fig. 8.**
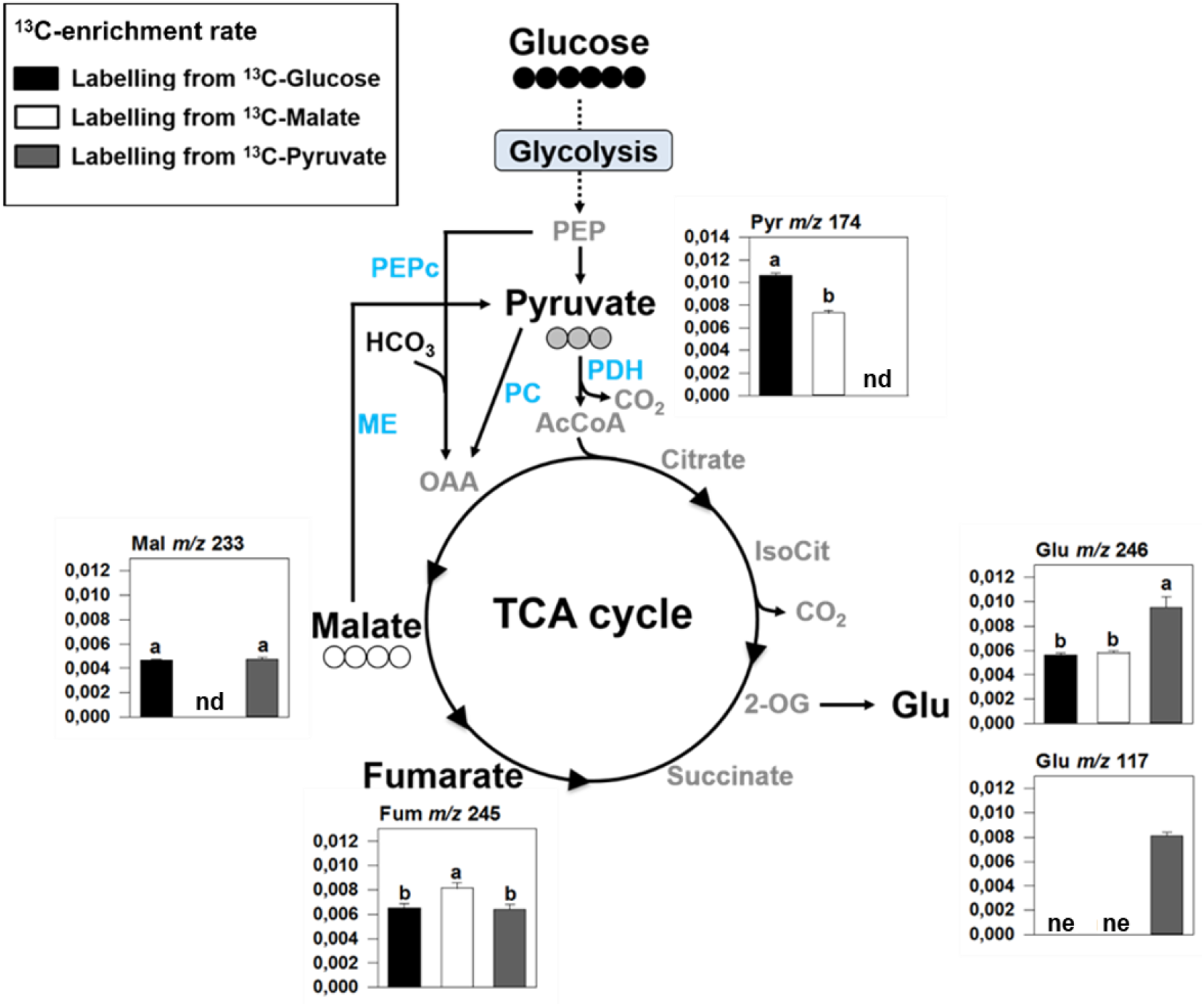
Diagram of TCA cycle and related metabolites showing the ^13^C-enrichment rate of mesophyll cells (MCs) from Arabidopsis leaves (MC-C3) fed with ^13^C-Glc, ^13^C-Mal or ^13^C-Pyr for 240 minutes, under light condition. The ^13^C-enrichment rate was determined by dividing the total relative ^13^C-enrichment by the ^13^C-feeding time, in minutes. Values are presented as mean ± SE (n=6). Metabolite fragments that were not detected by GC-MS in a specific labelling substrate were identified as not analysed (nd). Metabolite fragments with absent ^13^C-enrichment were identified as no ^13^C-enrichment observed (ne.). Different letters indicate significant difference among labelling substrates by Tukey’s test (*P* < 0.05). Abbreviations: Metabolites (AcCoA, acetyl-coenzyme A; Fum, fumarate; Glc, glucose; Glu, glutamate; IsoCit, isocitrate; Mal, malate; OAA, oxaloacetate; PEP, phospho*enol*pyruvate; Pyr, pyruvate; 2-OG, 2-oxoglutarate); Enzymes (ME, malic enzyme; PC, pyruvate carboxylase; PDH, pyruvate dehydrogenase; PEPc, phospho*enol*pyruvate carboxylase).

We next analysed Glu *m/z* 117 in MC-C3 in plants impaired in the thioredoxin system using a *trxo1* mutant and a *ntra ntrb* double mutant. TRX *o1* is localized in the mitochondrial matrix, while NTRA and NTRB are dually localized in the mitochondrial matrix and in the cytosol. Following provision of ^13^C-Mal, no ^13^C-enrichment in Glu *m/z* 117 was observed. Following provision of ^13^C-Pyr, we observed ^13^C-enrichment in all genotypes but without any differences in labelling. However, provision of ^13^C-Glc resulted in ^13^C-enrichment in Glu *m/z* 117 exclusively in the mutants, suggesting that a flux route that is completely inactive in the WT is activated in the mutants (Figure **9a**). More specifically, the impairment of the NTR/TRX system favours PEPc-derived ^13^C incorporation into the carbon 1 of the Glu backbone (Fig. **9b**). Since the impact was similar in both mutant lines, this adjustment is clearly caused by impaired regulation inside the mitochondrial and direct TRX *o1*-mediated regulation of cytosolic PEPc is unlikely. Instead, de-repression of a matrix enzyme, such as PDH, provides a plausible account for the observed re-routing of flux (Fig. **10**).

**Fig. 9.**
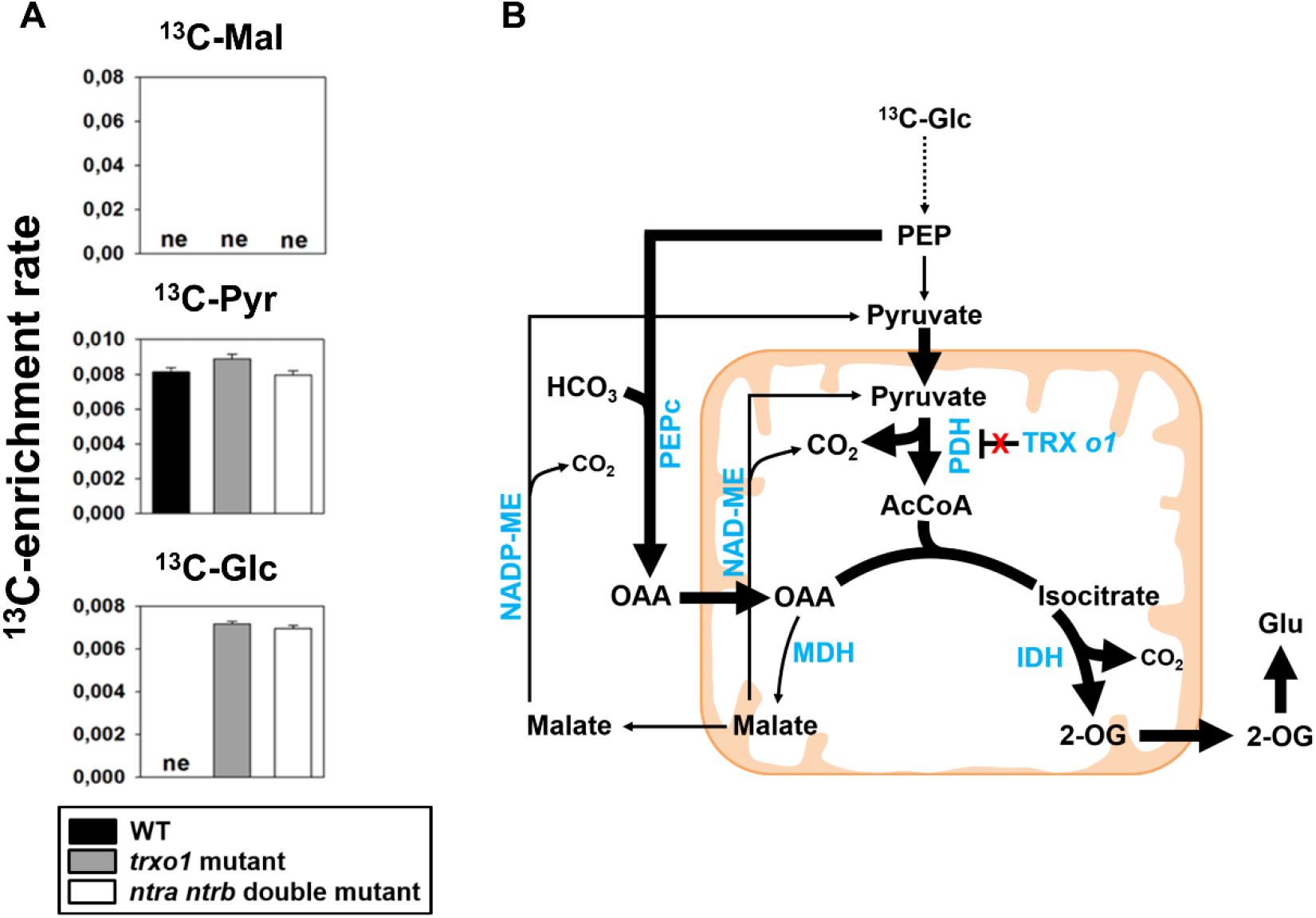
A) ^13^C-enrichment rate into Glu *m/z* 117 from mesophyll cells (MCs) of Arabidopsis leaves of wild-type (WT), *trxo*1 mutant and *ntra ntrb* double mutant fed with ^13^C-Mal, ^13^C-Pyr or ^13^C-Glc for 240 minutes in the light. The ^13^C-enrichment rate was determined by dividing the total relative ^13^C-enrichment by the ^13^C-feeding time, in minutes. No ^13^C-enrichment in Glu *m/z* 117 is identified as no ^13^C-enrichment observed (ne). Values are presented as mean ± SE (n=6). **B)** Schematic representation of the differential TCA-cycle metabolic fluxes in *trxo*1 mutant. In this scenario, the lack of TRX *o1* lifts the inhibition upon PDH, increasing the carbon fluxes toward Glu synthesis from glycolysis and PEPc activity, evidencing the importance of the redox regulation mediated by this TRX in modulating metabolic fluxes throughout the TCA cycle and associated pathways in illuminated MCs. Thicker arrows indicate higher carbon fluxes. Abbreviations: Metabolites: AcCoA, acetyl-coenzyme A; Glc, glucose; Glu, glutamate; Mal, malate; OAA, oxaloacetate; PEP, phospho*enol*pyruvate; Pyr, pyruvate; 2-OG, 2-oxoglutarate. Enzymes, highlighted in light blue: IDH, isocitrate dehydrogenase; MDH, malate dehydrogenase; NAD-ME, NAD-malic enzyme; NADP-ME, NADP-malic enzyme; PDH, pyruvate dehydrogenase; PEPc, phospho*enol*pyruvate carboxylase.

**Fig. 10.**
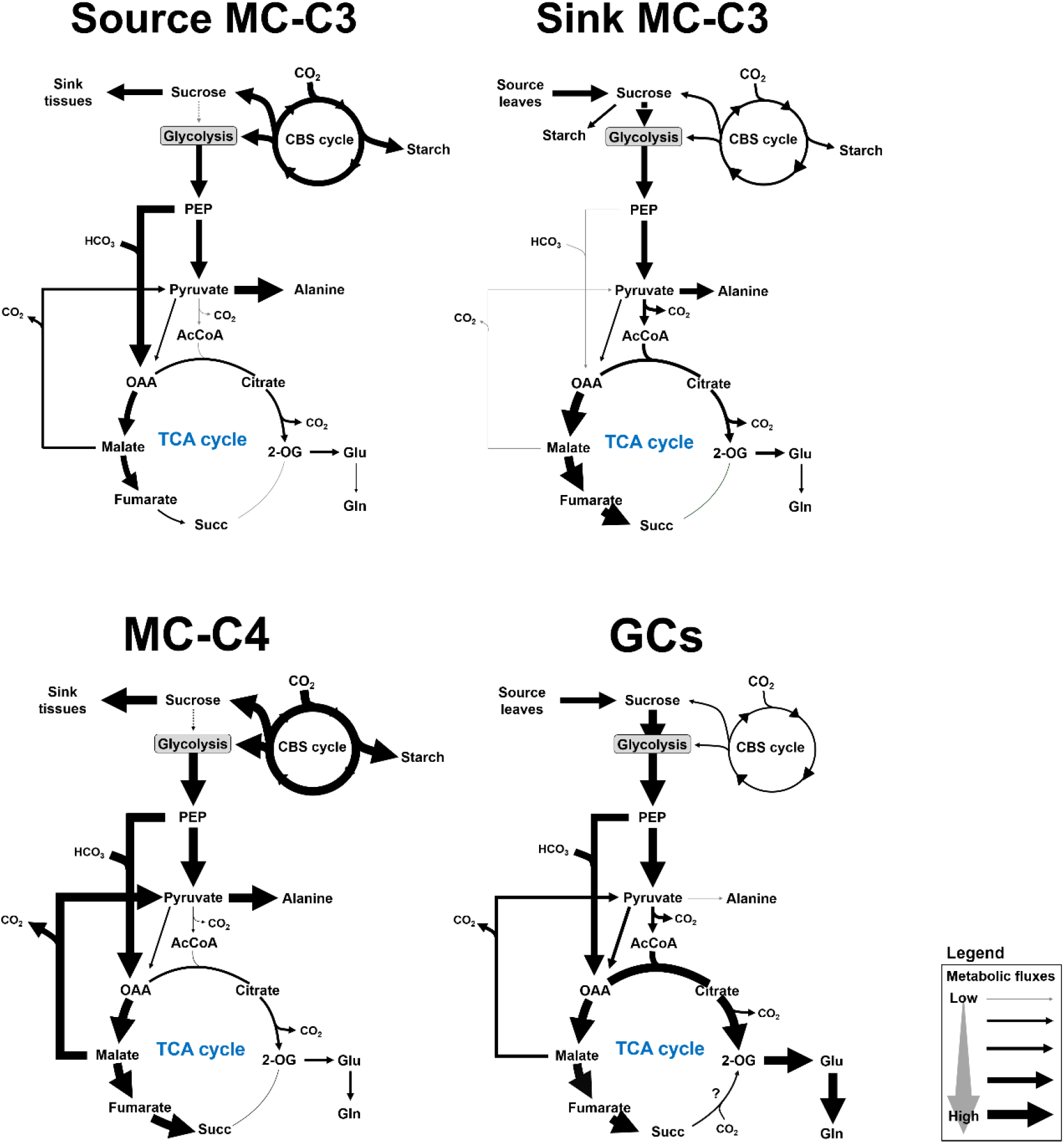
Schematic representation of the ^13^C distribution throughout the TCA cycle and associated pathways in mesophyll cells of C3 (MC-C3) and C4 (MC-C4) plants and in guard cells (GCs) of C3 plants. The ^13^C distribution over the TCA cycle suggests that GCs have a particular metabolic flux mode, with similarities with sink MC-C3. For instance, sink MC-C3 has a higher carbon flux toward Glu synthesis than source MC-C3, which resembles GCs. Moreover, sink MC-C3 and GCs show higher ^13^C-flux through the C4-branch of the TCA cycle, whilst source MC-C3 has highly restricted glycolytic carbon flux toward the TCA cycle. However, it seems likely that the GCs have higher fluxes through the C6 branch of the TCA cycle, when compared to all MCs. GCs also differ from MCs, in which lower glycolytic flux toward Ala synthesis in GCs was observed. Abbreviations: Metabolites: AcCoA, acetyl-coenzyme A; Glu, glutamate; Gln, glutamine; OAA, oxaloacetate; PEP, phospho*enol*pyruvate; Succ, succinate; 2-OG, 2-oxoglutarate. Others: CBS cycle, Calvin-Benson-Bassham cycle.

## Discussion

### Feasibility and potential of GCMS-based ^13^C-positional isotopomer labelling

The discrimination of the fate of ^13^C released from a labelled substrate and the identification of the precise atom position of the ^13^C incorporation in a metabolite are the major challenges in MFA. This is particularly true for plants due to both the intrinsic complexity of plant cell metabolism and limitations of available techniques for metabolomics (Sweetlove & Fernie, 2013). Despite all the constraints, it is currently feasible to identify ^13^C-labelling down to the level of specific positions in a molecule and ^13^C-NMR has proven particularly powerful for ^13^C-MFA (Zamboni *et al*., 2009). However, NMR provides less sensitivity than MS. Hence, NMR has not been used for guard cell MFA so far very likely due to the very large quantity of guard cells that would be required for such an analysis (Medeiros *et al*., 2019). An alternative to overcome this limitation and to improve MFA is the determination of positional isotope labelling in compounds identified by MS (Choi *et al*., 2012). GC-EI-MS is particularly suitable for the establishment of such an approach, given that it offers high resolution/sensibility with a stable fragmentation pattern due to the strong energy used in EI ionization. However, the lack of a clear fragmentation map has restricted the use of data from this platform in MFA approaches.

Aiming to overcome the technical limitations of MS-based MFA, we established a specific fragmentation map of TCA cycle-related metabolites that allowed us to investigate the ^13^C distribution from PEPc activity into Mal, Glu and Gln using data provided by MS-based studies. We were able to investigate the contribution of glycolysis, the TCA cycle and PEPc activity for the synthesis of Glu and Gln. The results obtained from our analysis are in overall agreement with recent ^13^C-NMR-based MFA (Abadie *et al*., 2017, 2018). The feasibility of our approach is supported by the fact that Mal 3TMS that are labelled specifically at the carbon 4 shows a shift from *m/z* 117 to *m/z* 118 when injected solely in a GC-EI-MS (Okahashi *et al*., 2019). Here, we demonstrate that the relative total ^13^C-enrichment in Mal *m/z* 117 is linearly correlated with the relative total ^13^C-enrichment in the carbon 4 of Mal. Following the intensity of the fragments *m/z* 117 and *m/z* 118, we could obtain insights into the ^13^C incorporation into Mal, which indirectly refers to the *in vivo* PEPc activity. As expected, our analysis showed that the ^13^C-enrichment in Mal *m/z* 118 and the ^13^C-enrichment in the carbon 4 of Mal is higher in MC-C4 than in MC-C3, evidencing a higher PEPc-mediated ^13^C incorporation into Mal in C4 when compared to C3 cells. This result highlights the effectiveness of the ^13^C-positional isotope labelling approach, which can be further applied to establish a method for PEPc activity measurements *in vivo* using GC-MS data. An alternative approach for the determination of PEPc activity *in vivo* has been recently established by ^13^C-NMR MFA (Abadie & Tcherkez, 2019). Here, we provide proof-of-concept for a more straightforward approach through the analysis of GC-EI-MS-based ^13^C-MFA. It is important to highlight that this approach is not limited to plant cells, given that GC-MS is one of the most common platforms used in metabolomics (Garcia & Barbas, 2011), offering a possibility to improve the resolution of metabolic flux maps across organisms. However, it is also important to highlight that our GC-APCI-MS indicates that a careful validation using positionally ^13^C-labelled standards and different MS platforms is recommended to avoid misleading in the flux maps reconstruction and interpretation.

### Non-cyclic metabolic flux modes of guard cell TCA cycle in the light

The results obtained here are in line with the idea that the TCA cycle of GCs shows different non-cyclic flux modes, in which the C4 and the C6-branch operate depending on substrate availability. Whilst the ^13^C derived from Suc degradation was strongly incorporated into Glu and Gln, suggesting the activation of the TCA cycle-C6 branch, the data obtained for GCs following provision of ^13^C-HCO_3_ show a higher ^13^C incorporation rate into Fum, when compared to MC-C3 following provision of ^13^CO_2_, indicating that GCs TCA cycle-C4 branch is also rapidly activated in the light. Whilst the fate of Gln, and to which extent and how it contributes to regulate stomatal opening are still open questions, the activation of the TCA cycle-C4 branch of GCs is an important mechanism to support stomatal opening by contributing to the osmotic balance of the cell and/or by stimulating ATP synthesis (Lawson & Matthews, 2020).

The non-cyclic flux mode of the TCA cycle separated in two branches is commonly observed in illuminated leaves (Tcherkez *et al*., 2012). Additional non-cyclic flux modes are found in microorganisms (Zhang & Bryant, 2011; Steinhauser *et al*., 2012). For instance, certain bacteria show a completely reverse TCA cycle as an important pathway for CO_2_ fixation (Evans *et al*., 1966; Buchanan & Arnon, 1990; Mall *et al*., 2018). In this context, it is interesting to note that in GCs following provision of ^13^C-HCO_3_, Succ content significantly increased from 0 (dark) to 24 min after light imposition and decreased substantially afterwards. Similarly, the total ^13^C enrichment in Succ *m/z* 247 significantly increased on the first 32 min upon light and then decreased towards 40 min. These results indicate that the carbon from ^13^C-HCO_3_ is used to the synthesis of Succ and that this metabolite is metabolized over time after a brief exposure to light. Intriguingly, the total relative ^13^C-enrichment in Glu *m/z* 246 showed a rapid increase from 32 to 40 min in GCs following provision of ^13^C-HCO_3_, in parallel to an increase in the intensity of both Glu M3 and M4. Glu *m/z* 246 contains four carbons of the Glu backbone, in which two carbons are derived from PDH activity and the other two from the TCA cycle. Although we did not detect Glu *m/z* 117-118 in this experiment, the results from ^13^C-Suc showed an increase in Glu *m/z* 118 after 30 min in the light. It seems likely therefore that all carbon atoms of Glu are labelled in GCs during dark-to-light transition by a combination of ^13^C derived from PEPc, glycolysis and the TCA cycle. Assuming that the TCA cycle does not work in a cyclic mode in the light, a reasonable hypothesis to explain the labelling in all carbon atoms of Glu in GCs following provision of ^13^C-HCO_3_ will be an association of Succ degradation and Glu synthesis. In this scenario, the TCA cycle would run in a non-cyclic, partially reverse mode. In terms of functionality for guard cell metabolism, this partial reverse mode would contribute to the overall CO_2_ fixation of the cell, given that a molecule of CO_2_ would be fixed from succinyl CoA to 2-OG (Buchanan & Arnon, 1990). However, the ^13^C-enrichment in all carbon atoms of Glu *m/z* 246 could also be interpreted by a combination of labelled carbons entering the TCA cycle through the activity of PDH, PC and PEPc using fully labelled Pyr and PEP and producing fully labelled OAA and AcCoA. In this scenario, the reduction observed on Succ labelling and content could be interpreted as a potential consumption of this metabolite by oxidative phosphorylation system. Further ^13^C-feeding experiments and enzymatic characterization of TCA cycle enzymes are thus necessary to test the hypothesis that guard cell TCA cycle runs in a non-cyclic, partially reverse mode to support Gln synthesis in the light.

### Similarities and differences between guard cells and sink-C3 mesophyll cells

GCs share several metabolic characteristics with sink MCs (Daloso *et al*., 2016a). In agreement with this concept, we found that the distribution pattern of the carbon derived from ^13^C-Suc to Gln in GCs is more related to sink rather than to source MCs. For instance, sink MC-C3 showed 3-fold higher total relative ^13^C-enrichment in Gln than source MC-C3. Similarly, GCs displayed high ^13^C enrichment rate in Gln from ^13^C-Suc, which was greater than MC-C3 or MC-C4 following provision of ^13^CO_2_. By contrast, a remarkable difference found between GCs and MCs was the ^13^C incorporation pattern into Ala. GCs following provision of ^13^C-Suc displayed a higher rate of ^13^C-enrichment in Pyr and much lower ^13^C-enrichment rate in Ala, as compared to MC-C3 following provision of ^13^CO_2_. Furthermore, whilst Pyr content reduced substantially over time, Ala content did not change in GCs following provision of ^13^C-Suc. This indicates that, during dark-to-light transition, glycolysis has a minor importance in providing carbon for the synthesis of Ala in GCs.

MC-C4 had an extremely rapid ^13^C enrichment in Ala, which is probably explained by a combination of labelled carbon coming from glycolysis and the activities of ME and AlaAT. Interestingly, GCs also showed an increase in the relative abundance of the isotopologue Pyr *m/z* 177 following provision of ^13^C-Suc over time, leading to statistically higher ^13^C-enrichment rate than MC-C3 following provision of ^13^CO_2_. This result suggests that the glycolytic flux toward Pyr is higher in GCs than MC-C3 in the light. Despite the lower ^13^C-incorporation into Ala, we observed a high incorporation of ^13^C derived from Pyr into Glu/Gln in GCs, as evidenced by the progressive increase in Glu *m/z* 248 over time. Furthermore, whilst Glu *m/z* 246 and Gln *m/z* 245 clearly had the incorporation of four labelled carbons in GCs following provision of ^13^C-Suc, only two labelled carbons were incorporated into Glu in sink MC-C3 following provision of ^13^C-Suc (Dethloff *et al*., 2017) and in MC-C3 following provision of ^13^CO_2_ (Fig. **S12**). Additionally, no ^13^C-enrichment in Gln *m/z* 245 was observed in MC-C3 and MC-C4 following provision of ^13^CO_2_. These results suggest that GCs have a higher metabolic flux toward Glu and Gln in the light, with preference for Gln synthesis, and that PDH offers no, or at least lesser, restriction to the entrance of carbon into the TCA cycle in GCs upon light, when compared to MCs. These results further evidence that GCs share metabolic characteristics with sink cells in terms of a high allocation of carbon to Gln synthesis, but with differential contributions from glycolysis and the TCA cycle.

### Glycolytic flux toward glutamate synthesis is repressed by the mitochondrial TRX system in mesophyll cells in the light

Our analysis reinforces the idea that the light inhibition of the mitochondrial PDH is an important step that modulates the metabolic fluxes from PEPc and glycolysis toward Glu synthesis. This idea is supported by the result in which Glu *m/z* 117 was only labelled following provision of ^13^C-Pyr, and not following provision of ^13^C-Mal and ^13^C-Glc. Although Glu *m/z* 246 was labelled by all substrates, it was labelled to a greater extent following provision of ^13^C-Pyr, which suggest the intensification of PDH activity by increased substrate availability. Given that Glu *m/z* 117 contains the carbon derived from PEPc fixation, it seems likely therefore that PEPc frequently uses the CO_2_ released by PDH and/or IDH. Furthermore, ^13^C-enrichment in Glu *m/z* 117 following provision of ^13^C-Glc exclusively in *trxo1* and *ntra ntrb* mutants suggests that the mitochondrial TRX system is a negative regulator of this pathway *in vivo*. In this scenario, the lack of TRX *o1* or both NTRA/NTRB lifts the inhibition upon PDH and/or IDH activity, increasing therefore the rate of labelled CO_2_ released from mitochondria and fixed by PEPc. However, mitochondrial IDH is known to be activated by TRXs *in vitro* (Yoshida & Hisabori, 2014) and the lack of *trxo1* did not change the activity of IDH in mitochondrial extracts (Daloso *et al*., 2015b). In contrast, it has been recently shown that both TRX *o1* and TRX *h2* inhibit the activity of the mitochondrial dihydrolipoamide dehydrogenase (mtLPD) *in vitro* (Reinholdt *et al*., 2019; Fonseca-Pereira *et al*., 2020), which correspond to the E3 subunit of PDH, which is indeed believed to be redox regulated (Balmer *et al*., 2004; Nietzel *et al*., 2020). Thus, it is possible that the ^13^C-enrichment in Glu *m/z* 117 in the absence of TRX *o1* or of both NTRA and NTRB would be mainly related to an increase in PDH activity. Alternatively, given that mitochondrial MDH is not redox regulated (Yoshida & Hisabori, 2016), another possibility is that TRX/NTR mutants have higher export of Mal to cytosol, in which its decarboxylation by ME could increase the amount of CO_2_ for PEPc. Taken together, these results provide further evidence that mitochondrial TRX/NTR system is an important control point to modulate metabolic fluxes in matrix carbon metabolism and beyond.

## Acknowledgments

This work was made possible through financial support from the National Council for Scientific and Technological Development (CNPq, Grant 428192/2018-1). J.K. and A.E. acknowledge funding of the German science foundation (DFG) grant (KO 2329/7-1) that is part of the SCyCode research consortium (FOR2816). We also thank the research fellowship granted by CNPq to D.M.D and the scholarships granted by the Brazilian Federal Agency for Support and Evaluation of Graduate Education (CAPES-Brazil) to A.G.D. and V.F.L. We thank the authors who kindly provided their mass spectral data from published work.

## List of author contributions

Data analysis was performed by AGD, VFL, AE, LA and DMD. Validation by analysing positionally labelled standards was carried out by AE, with the supervision of JK. All authors contributed to the establishment of the method and to write the final manuscript; DMD obtained funding and is responsible for this article.

## Supplemental information

Additional Supporting Information may be found in the online version of this article.

**Fig. S1** Estimated EI-fragmentation map of TMS-derivatized metabolites of, or related to, the TCA cycle

**Fig. S2** Analysis of glutamate (Glu) standards via GC-EI-TOF-MS

**Fig. S3** Mass spectral analysis of malate standards

**Fig. S4** Detailed mass spectral analyses of the malate fragment *m/z* 117

**Fig. S5** Relationship between the mathematical estimation of the ^13^C-enrichment in the carbon 4 of malate (Mal) with the relative total ^13^C-enrichment in the fragment *m/z* 117 of Mal

**Fig. S6** Mass spectral analysis of aspartate standards

**Fig. S7** Relative metabolite content of Pyr, Asp and Ala

**Fig. S8** Relative abundance of mass isotopomers of Pyr, Asp and Ala

**Fig. S9** Relative abundance of mass isotopomers of Fum, Mal and Succ

**Fig. S10** Relative metabolite content of Fum, Mal and Succ

**Fig. S11** Relative total ^13^C-enrichment in the carbon 4 of malate

**Fig. S12** Relative abundance of mass isotopomers of Fum, Mal and Succ

**Fig. S13** Relative metabolite content of Glu and Gln

**Fig. S14** Relative metabolite content and total relative ^13^C-enrichment in glutamate and succinate in guard cells fed with ^13^C-HCO_3_

**Table S1** General description of the previous metabolic flux studies used for the multi-species/cell-types analysis.

## References

Abadie C, Bathellier C, Tcherkez G. 2018. Carbon allocation to major metabolites in illuminated leaves is not just proportional to photosynthesis when gaseous conditions (CO_2_ and O_2_) vary. New Phytologist 218: 94–106.

Abadie C, Lothier J, Boex-Fontvieille E, Carroll A, Tcherkez G. 2017. Direct assessment of the metabolic origin of carbon atoms in glutamate from illuminated leaves using ^13^C-NMR. New Phytologist 216: 1079–1089.

Abadie C, Tcherkez G. 2019. In vivo phospho enol pyruvate carboxylase activity is controlled by CO_2_ and O_2_ mole fractions and represents a major flux at high photorespiration rates. New Phytologist 221: 1843–1852.

Antoniewicz MR. 2013. Tandem mass spectrometry for measuring stable-isotope labeling. Current Opinion in Biotechnology 24: 48–53.

Antoniewicz MR, Kelleher JK, Stephanopoulos G. 2007. Accurate assessment of amino acid mass isotopomer distributions for metabolic flux analysis. Analytical Chemistry 79: 7554–7559.

Araújo WL, Nunes-Nesi A, Osorio S, Usadel B, Fuentes D, Nagy R, Balbo I, Lehmann M, Studart-Witkowski C, Tohge T, et al. 2011. Antisense Inhibition of the Iron-Sulphur Subunit of Succinate Dehydrogenase Enhances Photosynthesis and Growth in Tomato via an Organic Acid–Mediated Effect on Stomatal Aperture. The Plant Cell 23: 600–627.

Araujo WL, Nunes-Nesi A, Trenkamp S, Bunik VI, Fernie AR. 2008. Inhibition of 2-Oxoglutarate Dehydrogenase in Potato Tuber Suggests the Enzyme Is Limiting for Respiration and Confirms Its Importance in Nitrogen Assimilation. Plant Physiology 148: 1782–1796.

Arrivault S, Obata T, Szecówka M, Mengin V, Guenther M, Hoehne M, Fernie AR, Stitt M. 2017. Metabolite pools and carbon flow during C4 photosynthesis in maize: 13 CO2 labeling kinetics and cell type fractionation. Journal of Experimental Botany 68: 283–298.

Balmer Y, Vensel WH, Tanaka CK, Hurkman WJ, Gelhaye E, Rouhier N, Jacquot J-P, Manieri W, Schurmann P, Droux M, et al. 2004. Thioredoxin links redox to the regulation of fundamental processes of plant mitochondria. Proceedings of the National Academy of Sciences 101: 2642–2647.

Beylot M, Previs SF, David F, Brunengraber H. 1993. Determination of the 13C-labeling pattern of glucose by gas chromatography-mass spectrometry. Analytical Biochemistry 212: 526–531.

Buchanan BB, Arnon DI. 1990. A reverse KREBS cycle in photosynthesis: consensus at last. Photosynthesis Research 24: 47–53.

Cheung CYM, Poolman MG, Fell DA, Ratcliffe RG, Sweetlove LJ. 2014. A Diel Flux Balance Model Captures Interactions between Light and Dark Metabolism during Day-Night Cycles in C3 and Crassulacean Acid Metabolism Leaves. Plant Physiology 165: 917–929.

Choi J, Grossbach MT, Antoniewicz MR. 2012. Measuring complete isotopomer distribution of aspartate using gas chromatography/tandem mass spectrometry. Analytical Chemistry 84: 4628–4632.

Daloso DM, dos Anjos L, Fernie AR. 2016a. Roles of sucrose in guard cell regulation. New Phytologist 211: 809–818.

Daloso DM, Antunes WC, Pinheiro DP, Waquim JP, Araújo WL, Loureiro ME, Fernie AR, Williams TCR. 2015a. Tobacco guard cells fix CO_2_ by both Rubisco and PEPcase while sucrose acts as a substrate during light-induced stomatal opening. Plant Cell and Environment 38: 2353–2371.

Daloso DM, Medeiros DB, dos Anjos L, Yoshida T, Araújo WL, Fernie AR. 2017. Metabolism within the specialized guard cells of plants. New Phytologist 216: 1018– 1033.

Daloso DM, Müller K, Obata T, Florian A, Tohge T, Bottcher A, Riondet C, Bariat L, Carrari F, Nunes-Nesi A, et al. 2015b. Thioredoxin, a master regulator of the tricarboxylic acid cycle in plant mitochondria. Proceedings of the National Academy of Sciences of the United States of America 112: E1392–E1400.

Daloso DM, Williams TCR, Antunes WC, Pinheiro DP, Müller C, Loureiro ME, Fernie AR. 2016b. Guard cell-specific upregulation of sucrose synthase 3 reveals that the role of sucrose in stomatal function is primarily energetic. New Phytologist 209: 1470–1483.

Dethloff F, Orf I, Kopka J. 2017. Rapid in situ 13C tracing of sucrose utilization in Arabidopsis sink and source leaves. Plant Methods 13: 1–19.

Dunnett CW. 1955. A Multiple Comparison Procedure for Comparing Several Treatments with a Control. Journal of the American Statistical Association 50: 1096– 1121.

Erban A, Martinez-Seidel F, Rajarathinam Y, Dethloff F, Orf I, Fehrle I, Alpers J, Beine-Golovchuk O, Kopka J. 2020. Multiplexed Profiling and Data Processing Methods to Identify Temperature-Regulated Using Gas Chromatography Coupled to Mass Spectrometry. In: Hincha DK, Zuther E, eds. Plant Cold Acclimation. New York, NY: Humana Press, 203–239.

Evans MC, Buchanan BB, Arnon DI. 1966. A new ferredoxin-dependent carbon reduction cycle in a photosynthetic bacterium. Proceedings of the National Academy of Sciences of the United States of America 55: 928–934.

Florez-Sarasa I, Obata T, Del-Saz NF, Reichheld JP, Meyer EH, Rodriguez-Concepcion M, Ribas-Carbo M, Fernie AR. 2019. The Lack of Mitochondrial Thioredoxin TRXo1 Affects in Vivo Alternative Oxidase Activity and Carbon Metabolism under Different Light Conditions. Plant and Cell Physiology 60: 2369– 2381.

Fonseca-Pereira P, Souza PVL, Hou LY, Schwab S, Geigenberger P, Nunes-Nesi A, Timm S, Fernie AR, Thormählen I, Araújo WL, et al. 2020. Thioredoxin h2 contributes to the redox regulation of mitochondrial photorespiratory metabolism. Plant Cell and Environment 43: 188–208.

Franzisky BL, Geilfus CM, Romo-Pérez ML, Fehrle I, Erban A, Kopka J, Zörb C. 2020. Acclimatisation of guard cell metabolism to long-term salinity. Plant Cell and Environment: 1–15.

Garcia A, Barbas C. 2011. Gas chromatography-mass spectrometry (GC-MS)-based metabolomics. Methods in molecular biology (Clifton, N.J.) 708: 191–204.

Gauthier PPG, Bligny R, Gout E,Mahé A, Nogués S, Hodges M, Tcherkez GGB. 2010. In folio isotopic tracing demonstrates that nitrogen assimilation into glutamate is mostly independent from current CO_2_ assimilation in illuminated leaves of Brassica napus. New Phytologist 185: 988–999.

Gotow K, Taylor S, Zeiger E. 1988. Photosynthetic Carbon Fixation in Guard Cell Protoplasts of Vicia faba L. Plant Physiol 86: 700–705.

Heise R, Arrivault S, Szecowka M, Tohge T, Nunes-Nesi A, Stitt M, Nikoloski Z, Fernie AR. 2014. Flux profiling of photosynthetic carbon metabolism in intact plants. Nature Protocols 9: 1803–1824.

De Jongh DC, Radford T, Hribar JD, Hanessian S, Bieber M, Dawson G, Sweeley CC. 1969. Analysis of Trimethylsilyl Derivatives of Carbohydrates by Gas Chromatography and Mass Spectrometry. Journal of the American Chemical Society 91: 1728–1740.

Kopka J, Schmidt S, Dethloff F, Pade N, Berendt S, Schottkowski M, Martin N, Dühring U, Kuchmina E, Enke H, et al. 2017. Systems analysis of ethanol production in the genetically engineered cyanobacterium Synechococcus sp. PCC 7002. Biotechnology for Biofuels 10: 1–21.

Lawson T, Matthews J. 2020. Guard Cell Metabolism and Stomatal Function. Annual Review ofPlant Biology 71: 273–302.

Lawson T, Oxborough K, Morison JIL, Baker NR. 2003. The responses of guard and mesophyll cell photosynthesis to CO_2_, O_2_, light, and water stress in a range of species are similar. Journal of Experimental Botany 54: 1743–1752.

Leimer KR, Rice RH, Gehrke CW. 1977. Complete mass spectra of the per-trimethylsilylated amino acids. Journal of Chromatography A 141: 355–375.

Lima VF, de Souza LP, Williams TCR, Fernie AR, Daloso DM. 2018. Gas Chromatography–Mass Spectrometry-Based ^13^C-Labeling Studies in Plant Metabolomics. In: Plant Metabolomics. 47–58.

Mall A, Sobotta J, Huber C, Tschirner C, Kowarschik S, Bačnik K, Mergelsberg M, Boll M, Hügler M, Eisenreich W, et al. 2018. Reversibility of citrate synthase allows autotrophic growth of a thermophilic bacterium. Science 359: 563–567.

Medeiros DB, da Luz LM, de Oliveira HO, Araújo WL, Daloso DM, Fernie AR. 2019. Metabolomics for understanding stomatal movements. Theoretical and Experimental Plant Physiology 9: 91–102.

Medeiros DB, Perez Souza L, Antunes WC, Araújo WL, Daloso DM, Fernie AR. 2018. Sucrose breakdown within guard cells provides substrates for glycolysis and glutamine biosynthesis during light-induced stomatal opening. Plant Journal 94: 583– 594.

Møller IM, Igamberdiev AU, Bykova N V, Finkemeier I, Rasmusson AG, Schwarzländer M. 2020. Matrix Redox Physiology Governs the Regulation of Plant Mitochondrial Metabolism through Posttranslational Protein Modifications. The Plant Cell 32: 573–594.

Nietzel T, Mostertz J, Ruberti C, Née G, Fuchs P, Wagner S, Moseler A, Müller-Schüssele SJ, Benamar A, Poschet G, et al. 2020. Redox-mediated kick-start of mitochondrial energy metabolism drives resource-efficient seed germination. Proceedings of the National Academy of Sciences of the United States of America 117: 741–751.

Okahashi N, Kawana S, Iida J, Shimizu H, Matsuda F. 2019. Fragmentation of Dicarboxylic and Tricarboxylic Acids in the Krebs Cycle Using GC-EI-MS and GC-EI-MS/MS. Mass Spectrometry 8: A0073–A0073.

Petersson G. 1972. Mass spectrometry of hydroxy dicarboxylic acids as trimethylsilyl derivatives. Rearrangement fragmentations. Organic Mass Spectrometry 6: 565–576.

Ratcliffe RG, Shachar-Hill Y. 2006. Measuring multiple fluxes through plant metabolic networks. Plant Journal 45: 490–511.

Reinholdt O, Schwab S, Zhang Y, Reichheld J-P, Fernie AR, Hagemann M, Timm S. 2019. Redox-Regulation of Photorespiration through Mitochondrial Thioredoxin o1. Plant Physiology 181: 442–457.

Robaina-Estévez S, Daloso DM, Zhang Y, Fernie AR, Nikoloski Z. 2017. Resolving the central metabolism of Arabidopsis guard cells. Scientific Reports 7: 1–13.

Sevilla F, Mart C, Jime A. 2020. Thioredoxin Network in Plant Mitochondria: Cysteine S-Posttranslational Modi fi cations and Stress Conditions. 11: 1–20.

Sienkiewicz-Porzucek A, Sulpice R, Osorio S, Krahnert I, Leisse A, Urbanczyk-Wochniak E, Hodges M, Fernie AR, Nunes-Nesi A. 2010. Mild reductions in mitochondrial NAD-dependent isocitrate dehydrogenase activity result in altered nitrate assimilation and pigmentation but do not impact growth. Molecular Plant 3: 156–173.

Souza LP de, Fernie AR, Tohge T. 2018. Carbon Atomic Survey for Identification of Selected Metabolic Fluxes. In: António C, ed. Methods in Molecular Biology. Plant Metabolomics. New York, NY: Springer New York, 59–67.

Steinhauser D, Fernie AR, Araújo WL. 2012. Unusual cyanobacterial TCA cycles: Not broken just different. Trends in Plant Science 17: 503–509.

Sulpice R, Sienkiewicz-Porzucek A, Osorio S, Krahnert I, Stitt M, Fernie AR, Nunes-Nesi A. 2010. Mild reductions in cytosolic NADP-dependent isocitrate dehydrogenase activity result in lower amino acid contents and pigmentation without impacting growth. Amino Acids 39: 1055–1066.

Sussmilch FC, Roelfsema MRG, Hedrich R. 2019. On the origins of osmotically driven stomatal movements. New Phytologist 222: 84–90.

Sweetlove LJ, Beard KFM, Nunes-Nesi A, Fernie AR, Ratcliffe RG. 2010. Not just a circle: Flux modes in the plant TCA cycle. Trends in Plant Science 15: 462–470.

Sweetlove LJ, Fernie AR. 2013. The Spatial Organization of Metabolism Within the Plant Cell. Annual Review of Plant Biology 64: 723–746.

Szecowka M, Heise R, Tohge T, Nunes-Nesi A, Vosloh D, Huege J, Feil R, Lunn J, Nikoloski Z, Stitt M, et al. 2013. Metabolic Fluxes in an Illuminated Arabidopsis Rosette. The Plant Cell 25: 694–714.

Tcherkez G, Boex-Fontvieille E, Mahé A, Hodges M. 2012. Respiratory carbon fluxes in leaves. Current Opinion in Plant Biology 15: 308–314.

Tcherkez G, Cornic G, Bligny R, Gout E, Ghashghaie J. 2005. In Vivo Respiratory Metabolism of Illuminated Leaves. Plant Physiology 138: 1596–1606.

Tcherkez G, Mahé A, Gauthier P, Mauve C, Gout E, Bligny R, Cornic G, Hodges M. 2009. In folio respiratory fluxomics revealed by ^13^C isotopic labeling and H/D isotope effects highlight the noncyclic nature of the tricarboxylic acid ‘cycle’ in illuminated leaves. Plant Physiology 151: 620–630.

Tovar-Méndez A, Miernyk JA, Randall DD. 2003. Regulation of pyruvate dehydrogenase complex activity in plant cells. European Journal of Biochemistry 270: 1043–1049.

Willmer C, Fricker M. 1996. Stomata. London: Chapman & Hall.

Yoshida K, Hisabori T. 2014. Mitochondrial isocitrate dehydrogenase is inactivated upon oxidation and reactivated by thioredoxin-dependent reduction in Arabidopsis. Frontiers in Environmental Science 2: 1–7.

Yoshida K, Hisabori T. 2016. Adenine nucleotide-dependent and redox-independent control of mitochondrial malate dehydrogenase activity in Arabidopsis thaliana. Biochimica et Biophysica Acta (BBA) - Bioenergetics 1857: 810–818.

Zamboni N, Fendt S-M, Rühl M, Sauer U. 2009. 13C-based metabolic flux analysis. Nature Protocols 4: 878–892.

Zhang Y, Beard KFM, Swart C, Bergmann S, Krahnert I, Nikoloski Z, Graf A, George Ratcliffe R, Sweetlove LJ, Fernie AR, et al. 2017. Protein-protein interactions and metabolite channelling in the plant tricarboxylic acid cycle. Nature Communications 8.

Zhang S, Bryant DA. 2011. The tricarboxylic acid cycle in cyanobacteria. Science 334: 1551–1553.

Zhang Y, Swart C, Alseekh S, Scossa F, Jiang L, Obata T, Graf A, Fernie AR. 2018. The Extra-Pathway Interactome of the TCA Cycle: Expected and Unexpected Metabolic Interactions. Plant Physiology 177: 966–979.

